# Engineering antigen-driven co-stimulation and T helper cell activity into TCR-T cells with CD8-41BB fusion receptors enhances anti-tumor activity

**DOI:** 10.64898/2026.07.27.739613

**Authors:** Ivy Dutta, Justin Oh, Liz Cam, Allie Luther, Palak Sharma, Ishina Balwani, James Peter, Dai Liu, Ian C. Miller, James R. Bowen, Lekshmi Maya, Jingyu Peng, Eleni Stampouloglou, Qingzhan Zhang, Yoko Kosaka, Jesse L. Coy, Jessica S. Mulkey, Evan F. Lind, Eliana Ruggiero, Chiara Bonini, Laura Sepp-Lorenzino, Birgit C. Schultes, Aaron Prodeus

**Author notes:** **Corresponding Authors:** Aaron Prodeus, Intellia Therapeutics.

## Abstract

Adoptive cell therapy using tumor antigen-targeting T cell receptors (TCRs) offers a compelling approach to treat both hematological cancers and solid tumors due to broad antigen accessibility and the ability to target cancer-specific neoantigens. However, unlike clinically validated second generation CAR-T cells bearing built-in co-stimulatory signaling modules (i.e. 41BB or CD28), TCR-T cells receive little to no co-stimulation within most tumor microenvironments leading to attenuated cellular responses. Additionally, CD4+ TCR-T cells engineered to express HLA-Class I restricted TCRs possess minimal T-helper cell activity and thus do not effectively mobilize CD8+ TCR-T cells or host anti-tumor immune responses. To address these limitations, we used CRISPR-Cas9 to engineer TCR-T cells with targeted integration of chimeric CD8 constructs containing intracellular co-stimulatory domains. We found that expression of wild-type CD8αβ, but not CD8αα, could promote CD4+ T cell activities in HLA-Class I restricted TCR-T cells. However, this was insufficient to drive durable anti-tumor responses in challenging tumor mouse models when using a high-affinity WT1-directed TCR. To address this, several CD8 co-stimulatory fusion constructs containing CD28 or 41BB intracellular domains were designed and screened, identifying two CD8-41BB based chimeras that substantially increased TCR-T cell activity relative to wild-type CD8αβ. WT1-TCR-T cells co-expressing the CD8-41BB fusions demonstrated not only enhanced CD4+ activity including strong and polarized Th1-type cytokine secretion, but also enhanced the proliferation, cytokine release, and cytotoxicity of CD8+ CTLs. Remarkably, when combined with *TGFBR2* gene disruption, WT1-TCR-T cells co-expressing CD8-41BB receptors were able to completely regress established cell line-derived ovarian tumors, showed robust *in vivo* expansion and persistence, and provided long-term protection from tumor rechallenge. Importantly, the specificity profile of the WT1-TCR including its HLA-A*02:01 restriction and WT1 peptide recognition motif was preserved upon expression of CD8-41BB. To simplify cell engineering processes for clinical applications, we configured a homology directed repair (HDR) cassette to allow for efficient CRISPR-Cas9-based insertion of both the TCR and CD8-41BB transgenes in the *TRAC* locus in a single step with >80% efficiency. Lastly, the enhanced activity conferred by CD8-41BB expression was validated with a second clinically relevant TCR targeting PRAME, suggesting this platform can be a universal approach for enhancing the therapeutic potential of TCR-based cell therapies.

## 2 INTRODUCTION

Identification and use of tumor-specific T cell receptors (TCRs) to direct T cell mediated anti-tumor responses have shown therapeutic potential but have not yet achieved the clinical breakthroughs associated with curative responses achieved by second-generation chimeric antigen receptor (CAR)-T cells in hematological cancers^1,2^. Despite this, TCR-based therapeutics remain attractive due to (i) broader antigen availability by targeting either intracellular or extracellular antigens presented by MHCs, (ii) high specificity, enabling targeting of specific cancer-related neoantigens, and (iii) increased sensitivity to antigen expression, which may limit tumor escape due to heterogeneous target expression. Yet TCR-T cells have inherent limitations that constrain their therapeutic potential. One major shortcoming is lack of intrinsic T-helper cell activity since CD4+ cells engineered to express HLA-Class I-restricted TCRs are missing the secondary CD8-MHC contact point required for a high avidity interaction at the immune synapse^3,4^. Another limitation of TCR-T cell function is the lack of co-stimulatory ligands available within tumor microenvironments, particularly in solid tumors, leading to blunted T cell activation and susceptibility to T cell anergy and/or exhaustion, which can be exacerbated in the context of an immune-inhibitory microenvironment and lack of CD4+ T helper support^5–7^. In contrast, second-generation CAR-T cells possess high CD4+ Th1 type helper activity, and receive direct antigen driven co-stimulation conferred by built-in co-stimulatory 41BB or CD28 signaling domains. Notably, the inclusion of CD28 or 41BB signaling has been critical for the clinical success of second-generation CAR-T cells relative to first generation CAR-T cells, which exhibited attenuated cytokine release and limited cell expansion potential^8^.

There have been multiple strategies proposed to address these limitations to improve TCR-T cell potency. Several groups have demonstrated enhancement of HLA-Class I-bearing CD4+ T cells by overexpressing CD8αα or CD8αβ co-receptors using viral-based gene transfer of the endogenous sequences^9–11^. However, this strategy does not address limitations associated with lack of co-stimulatory activity. To this end, various approaches have emerged, including gene transfer of synthetic “switch receptors” containing an extracellular domain of an inhibitory receptor (ie. PD-1^12,13^, CD200R^14^, Fas^15^) fused with an intracellular stimulatory domain (CD28 or 41BB) to co-opt immunosuppressive signals. While promising, this strategy may be restrained in practice since the co-stimulatory activity is not antigen-driven and expression of inhibitory ligands is heterogeneous within tumors, across patients, and across tumor types^16–18^. Additionally, in isolation this approach will not address the lack of CD4+ T helper activity. Lenti- or retroviral-based methods to simultaneously transfer genes encoding the TCRαβ, CD8αβ, and switch receptors in the same cell poses significant logistical and biological challenges, including the need to manufacture multiple or large viral vectors and a potential genotoxicity risk related to semirandom integration events^19^.

To address these limitations, we leveraged CRISPR-Cas9 genome editing for precise insertion of tumor specific TCRs along with co-expression of constructs containing CD8-co-stimulatory chimeric receptors. This approach led to highly efficient site-specific integration of the TCR and CD8 transgenes into the *TRAC* or *AAVS1* safe harbor loci. In combination with a high-avidity WT1-TCR^20^, we screened several CD8-CD28 and CD8-41BB chimeric designs, identifying two candidates that showed robust enhancement of both CD4+ and CD8+ T cells *in vitro* and promoted powerful anti-tumor responses in challenging *in vivo* ovarian cancer and AML mouse models. Notably, the specificity profile of the WT1-TCR was preserved upon CD8-41BB co-receptor expression. The lead CD8-41BB receptors demonstrated similarly enhanced activity with a PRAME-specific TCR, indicating the broader utility of this approach for HLA-Class I-restricted TCRs.

## 3 Results

### 3.1 CRISPR-Cas9 based insertion of both TCRαβ and CD8αβ transgenes to elicit CD4+ T helper cell activity

In a previous study, we reported the identification of a specific WT1_37–45_-targeting TCR restricted to HLA-A*02:01 with applicability to both hematologic and solid tumors^20^. In contrast to WT1_126-134_-targeting TCRs that have been tested clinically in AML and require processing of the target epitope by the immunoproteasome^21^, the WT1^37–45^ peptide is broadly processed by the standard proteasome, thereby expanding the clinical applicability to include WT1-expressing solid tumors such as ovarian cancer^11,22,23^. Further, our previous study highlighted the benefit of using CRISPR-Cas9 to engineer T cells with highly efficient, targeted integration of this TCR into the *TRAC* locus (TCRα) combined with knockout of the endogenous TCRβ chain to maximize TCR expression and activity while minimizing TCR chain mispairing.

The WT1-TCR was found to be highly avid and capable of inducing degranulation even in CD4+ T cells in response to peptide-pulsed tumor cells. However, this required a 10-100-fold increase in antigen levels relative to CD8+ cells, a finding consistent with observations reported for other highly avid HLA-Class I-targeting TCRs^24^. To test the physiological relevance of this a 697 acute lymphocytic leukemia (ALL) tumor cell line expressing endogenous levels of WT1 was co-cultured with WT1-TCR T cells. Within these cultures, we observed minimal activation of CD4+ WT1-TCR cells, with no detectable proliferation or expression of IFN-γ and TNF-α **(Figure 1a)**, indicating minimal CD4+ T cell help is conferred in physiological settings where WT1 peptide presentation is limited. Similar results were observed when using other AML and ovarian cancer cell lines expressing endogenous levels of WT1 (see below), suggesting that effective CD4 T cell help engagement requires more than a high-avidity TCR.

**Figure 1.**
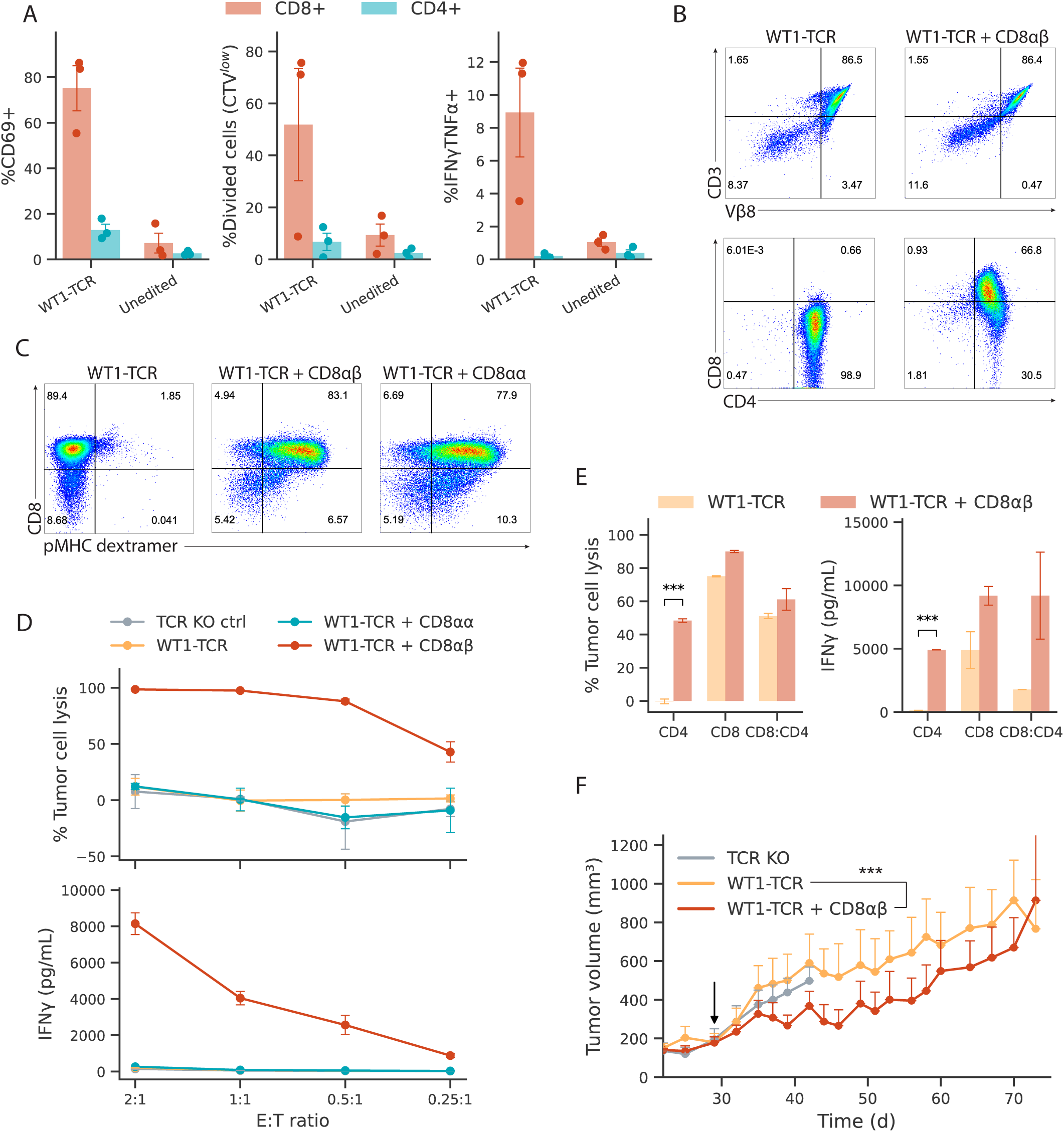
CRISPR-Cas9-mediated dual-targeted integration of CD8αβ and WT1-TCR enhances CD4+ T cell activity. **(a)** Flow cytometry was used to measure WT1-TCR T cell activation (CD69+), proliferation (Cell Trace Violet, CTV^low^), and cytokine expression (IFN-γ+ TNFα+) in CD4+ or CD8+ T cells after co-culture with 697 ALL cell line expressing endogenous levels of WT1 (n=3 donors, Mean ± SEM). Unedited cells from matching donors served as negative control. **(b)** CRISPR-Cas9-mediated dual targeted integration of the WT1-TCR (top) and CD8 co-receptors (bottom). Flow cytometric detection of WT1-TCR expression was achieved using an antibody specific for the T cell receptor beta variable gene segment (TRBV) used by this TCR (Vβ8). **(c)** Binding of fluorescently labelled WT1-dextramer to CD4+ WT1-TCR T cells expressing CD8αβ or CD8αα. **(d)** 697 ALL cytotoxicity (left) and IFN-γ secretion (right) from CD4+ WT1-TCR T cells expressing CD8αβ or CD8αα (n=2, Mean ± SD). **(e)** OVCAR3 cytotoxicity (left) and IFN-γ secretion (right) from WT1-CD4+, CD8+, or mixed populations of CD4+ and CD8+ T cells with or without CD8αβ insertion. **(f)** NOG-IL15 mice bearing ≥200mm^3^ OVCAR3 tumors were treated with 1×10^7^ T cells on day 29 (black arrow). Tumor volume was monitored over time, showing modest growth inhibition by WT1-TCR + CD8αβ T cells (n=6) compared to WT1-TCR T cells without CD8 expression (n=5) or TCR KO negative control cells (n=5). ***p<0.001.

Several groups have shown that overexpression of the CD8 co-receptor by lentiviral gene transfer can promote the effector activity of CD4+ T cells bearing HLA-Class I restricted TCRs^9–11^. To allow for site-specific insertion and high, uniform expression of TCR and CD8 transgenes, we built upon our previous methodology leveraging CRISPR-Cas9 for dual-targeted knock in (dual TKI) of the WT1-TCRαβ chains and CD8 chains into the *TRAC* and *AAVS1* loci, respectively, combined with knockout of the endogenous TCRβ. *AAVS1* was chosen as a suitable site for a second integration due to its nature as a “safe harbor” region and established precedent for transgene insertion^25,26^. The dual TKI process was highly efficient, achieving >80% insertion of the WT1-TCR and >60% insertion of the CD8 co-receptor, with both transgenes showing homogenous expression levels **(Figure 1b)**.

CD8 is most commonly found on T cells as an αβ heterodimer; however, an αα homodimer is expressed by a distinct population of lymphocytes and was reported to enhance the CD4+ activity of cells expressing an HLA-Class I-restricted MAGE-A4 specific TCR^9^. When expressed with the WT1-TCR, both CD8αα and CD8αβ restored the ability of the CD4+ T cells to bind a WT1-MHC dextramer **(Figure 1c)**. However, only CD8αβ expression conferred the ability of CD4+ WT1-TCR T cells to kill and secrete IFN-γ in response to 697 tumor cells **(Figure 1d)**. Further evaluation using fractionated CD4+, CD4+CD8+, or CD4-CD8+ cells **(Figure S1a)** showed that CD8αβ expression enhanced the lytic ability **(Figure S1b)** and cytokine release from CD4+WT1-TCR+ cells including increased secretion of IFN-γ, TNFα, and GM-CSF **(Figure S1c)**. Endogenous CD8+ (CD8+CD4-) cells with targeted insertion of CD8αβ had marginal increases in cytoxicity and secretion of GM-CSF, Granzyme B, and TNF-α, but not IFN-γ **(Figure S1c)**. Similarly, when challenged with a solid tumor ovarian cancer cell line (OVCAR3: HLA-A*02:01+, WT1+), WT1-TCR CD4+ T cells with CD8αβ insertion demonstrated increased cytotoxicity and IFN-γ secretion **(Figure 1e)**. Given the enhanced activity *in vitro,* we further evaluated this approach *in vivo* using an OVCAR3 mouse xenograft model. This model was previously found to be refractory to WT1-TCR T cell treatment when large subcutaneous tumors (≥200 mm^3^) were established prior to T cell dosing. Here, CD8αβ-inserted WT1-TCR T cells demonstrated significantly improved tumor control relative to the WT1-TCR only and TCR KO control groups. However, the responses were only partial and transient, with tumor control lapsing ∼25-days after treatment **(Figure 1f)**.

### 3.2 Screening of CD8 transgenes fused with cytoplasmic CD28 and 41BB co-stimulatory domains

To further enhance the potency of our WT1-TCR T cells, we hypothesized that including CD28 or 41BB intracellular domains in the CD8 transgene would be an elegant approach to provide co-stimulatory signals in an antigen (pMHC)-dependent manner, while still promoting CD4+ T cell activity. To this end, we designed and tested numerous constructs containing CD28 or 41BB signaling domains fused to either, or both, of the CD8α or CD8β chains following their respective transmembrane domains **(Figure 2a)**. Additionally, we designed control constructs with either deletion of the CD8α and/or CD8β intracellular domains. Bulk T cells engineered with dual TKI of the WT1-TCR and CD8 chimeras were assayed for IFN-γ secretion in co-cultures with OCI-AML3 tumor cells (WT1-low, HLA-A*02:01+). As anticipated, the CD8αβ construct enhanced IFN-γ secretion by ∼3-fold relative to WT1-TCR only cells **(Figure 2b)**, while the control construct with deletion of the CD8α and CD8β intracellular domains showed IFN-γ secretion comparable to that of the WT1-TCR only cells. Strikingly, two constructs containing the 41BB intracellular domain (ICD) on either the CD8α chain alone or both the CD8α and CD8β chains (hereafter referred to as CD8αβ-BBx1 or CD8αβ-BBx2, respectively) further enhanced IFN-γ secretion by ∼8.5-to 11-fold relative to WT1-TCR only cells. No other constructs, including those with the CD28 domain nor those with the 41BB domain added solely to the CD8β chain, showed improvement beyond wild-type CD8αβ. The finding that the 41BB ICD imparted increased activity upon replacing the CD8α ICD is notable as this is the domain responsible for mediating Lck signaling.

**Figure 2.**
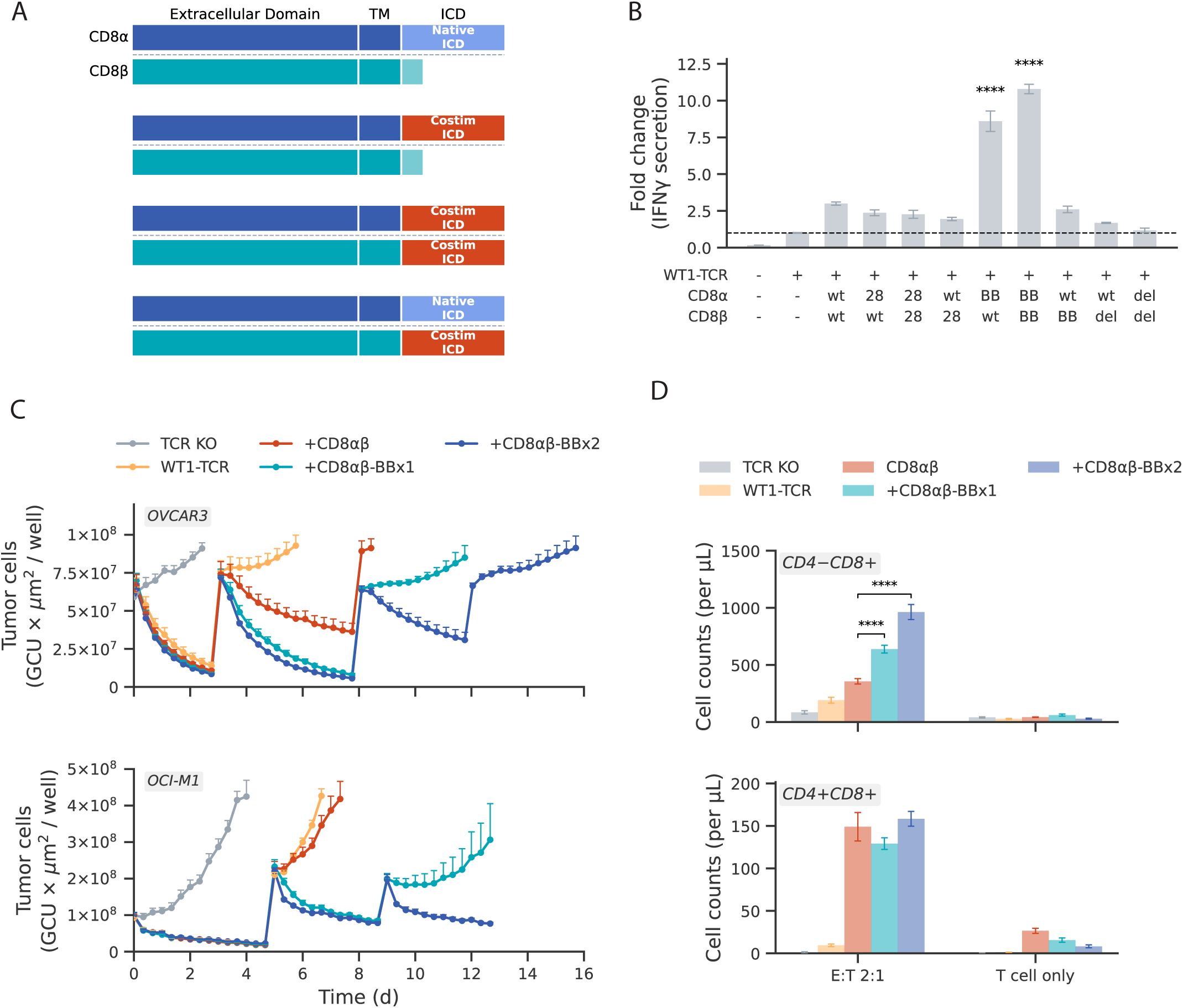
Design and screening of synthetic CD8 chimeras identified potent CD8-41BB co-stimulatory co-receptors. **(a)** Schematic of chimeric receptor designs tested, where the intracellular domains (ICDs) of CD8α, CD8β, or both, were replaced with CD28 or 41BB signaling domains. **(b)** Bulk T cells with dual targeted integration of the WT1-TCR and the indicated CD8α/CD8β chimeric sequences were co-cultured with OCI-AML3 tumor cells and IFN-γ secretion was measured in supernatants. Data were normalized to WT1-TCR only cells. Bars represent the mean ± SD (n=3). **(c)** WT1-TCR T cells expressing CD8αβ-BBx1 or CD8αβ-BBx2 show prolonged tumor killing in serial rechallenge assays with repetitive addition of OVCAR3 or OCI-M1 tumor cells. Symbols represent mean ± SD (n=3). **(d)** Proliferation of CTV-labelled WT1-TCR T cells expressing the indicated CD8 co-receptors after 4 days of co-culture with OCI-M1 tumor cells. Enhanced proliferation was observed in CD8+ T cells with CD8-41BB co-receptor insertion. Data represents CTV^low^WT1-TCR+ CD4-CD8+ or CD4+CD8+ cells quantified by counting beads (n=3, mean ± SD). T cell only groups served as baseline control. (n=2). ****p<0.001.

To assess the functional advantages provided by these two constructs beyond IFN-γ release, serial tumor rechallenge assays were performed using live cell imaging of T cell co-cultures with GFP-labelled OCI-M1 (AML, HLA-A*02:01+) or OVCAR3 cells. In these assays, WT1-TCR T cells expressing CD8αβ-BBx1 or -BBx2 chimeras markedly improved the long-term tumor cell killing capacity of WT1-TCR T cells relative to wild-type CD8αβ expression **(Figure 2c)**. Flow cytometry-based cell quantification of the OCI-M1/WT1-TCR T cell co-cultures showed enhanced proliferation in endogenous CD8+ (CD4-CD8+) T cells expressing CD8-41BB receptors (BBx2 > BBx1 > CD8αβ) **(Figure 2d)**. Among CD4+ cells, all three CD8 co-receptor groups had comparable levels of proliferation, while no cell division was observed in cells expressing only the WT1-TCR. No proliferation was observed in the absence of tumor cells in any of the conditions, confirming that the CD8-41BB receptor activity is strictly TCR-dependent.

Palmitoylation of transmembrane-adjacent motifs in both the CD8α and CD8β chains has been reported in both mouse and human CD8 receptors, however, the function of this post-translational modification is less clear. For mouse CD8, palmitoylation of the CD8β chain has been shown to promote recruitment of CD8αβ into lipid rafts^27^. For human CD8, the CD8α-chain palmitoylation domain along with the CD8β palmitoylation site and an adjacent arginine rich motif have all been implicated in lipid raft localization^28^. Our initial designs retained the CD8α-chain palmitoylation motif but deleted the HLCCRRRR motif encoded in the CD8β intracellular domain upon fusion with 41BB or CD28 **(Figure S2a)**. Reintroduction of these motifs did not increase membrane expression or activity of BBx2 designs, nor did it lead to previously unappreciated activity in fusions where a modified CD28 or 41BB ICD was added to the CD8β chain only **(Figure S2b,c)**. The dispensability of these motifs in the CD8β chain is consistent with findings that, for human CD8αβ, the self-assembly of the heterodimer mediated by extracellular regions, rather than palmitoylation, is sufficient for lipid raft localization^28^. Lastly, in an effort to retain Lck signaling, we engineered CD8 chimeras containing 41BB added to the end of the full length CD8α and/or CD8β chains **(Figure S2d)**. However, these constructs did not to demonstrate activity beyond that seen with wild-type CD8αβ **(Figure S2e)**. Thus, only the top performing CD8αβ-BBx1 and CD8αβ-BBx2 fusions were selected for further characterization.

### 3.3 CD8-41BB fusions augment activities of both CD4 and CD8 TCR-T cells

To explore the contributions of the CD8 chimeric receptors to CD4+ and CD8+ T cell activities independently, we separated CD4+ and CD8+ cells following CRISPR-Cas9 cell engineering and performed cytotoxicity and cytokine release assays using the OVCAR3 cell line. Short-term *in vitro* cytotoxicity **(Figure 3a)** measured over 3 days demonstrated that CD4+WT1-TCR+ cells expressing CD8αβ, CD8αβ-BBx1, and CD8αβ-BBx2 all imparted comparable cytotoxicity, with detectable killing observed at E:T ratios as low as a 1:8, while non-CD8 expressing CD4+WT1-TCR cells had only minimal toxicity at the highest E:T ratio tested (2:1). All CD8 constructs also modestly increased the cytotoxicity capacity of purified CD8+ T cells, demonstrated at the lowest E:T ratio (1:8). Overexpression of CD8αβ in CD8+ T cells has been previously documented to enhance TCR-T cell potency ^29^. This could be due to increased cell surface expression levels of CD8αβ **(Figure 3b)**, thereby increasing the avidity of the TCR-pMHC interaction.

**Figure 3.**
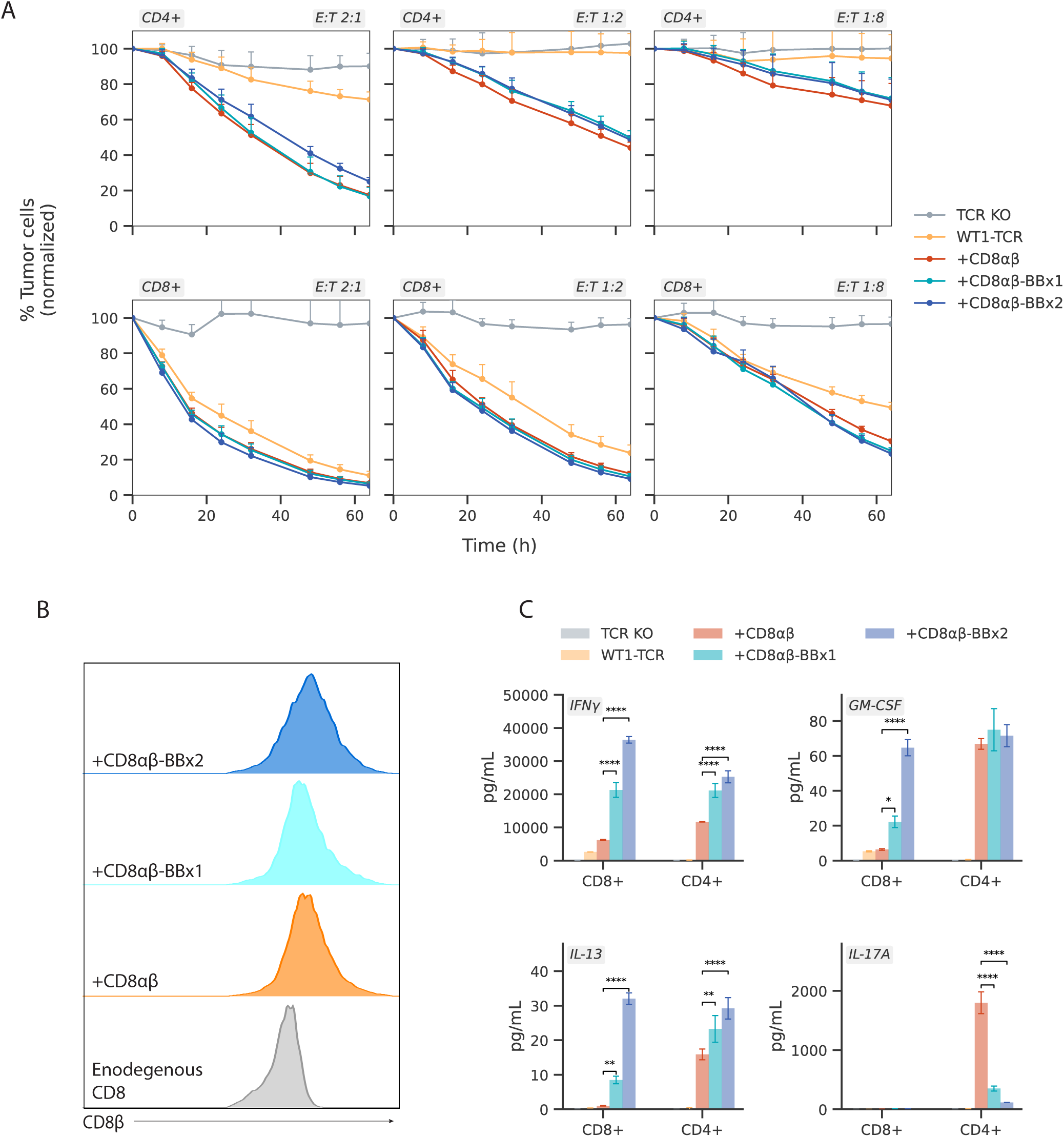
Contribution of CD8-41BB chimeras to CD4+ and CD8+ T cell function. **(a)** WT1-TCR+ CD4+ (top panels) and CD8+ (bottom panels) T cells were co-cultured with GFP+ OVCAR3 cells at the indicated E:T ratios. Enhanced cytotoxicity upon CD8 or CD8-41BB co-receptor expression is observed as GFP signal loss over time. Symbols represent mean ± SEM with experimental wells normalized to tumor cell only control wells. **(b)** Representative flow cytometric plots showing increased CD8 surface expression in endogenous CD8+(CD4-) T cells following CRISPR-Cas9-mediated insertion of the transgenic CD8 co-receptors. Data is representative of multiple independent studies. **(c)** Quantification of IFN-γ, GM-CSF, IL-13, and IL-17A secreted by WT1-TCR+ CD4+ or CD8+ T cells expressing the indicated CD8 co-receptor after 24h co-culture with OVCAR3 tumor cells. The introduction of 41BB signaling domains polarized T cell responses with increased IFN-γ and IL-13 secretion and reduced IL-17A secretion relative to the wild-type CD8αβ. Bars represent mean ± SD; data are representative of multiple independent studies. (*p<0.05, **p<0.01, ****p<0.001).

In contrast to the long-term killing studies with mixed CD4+ and CD8+ T cells, these short-term cultures did not reveal differences in killing activities across the CD8 receptors. However, analysis of IFN-γ, IL-17A, GM-CSF and IL-13 secretion showed differential cytokine profiles, with enhancement of Tc1 and Th1 activity in CD8-41BB-expressing cells **(Figure 3c)**. Specifically, CD8+ cells expressing CD8αβ-BBx1 or CD8αβ-BBx2 showed significantly increased IFN-γ secretion relative to CD8αβ. Additionally, the CD8-41BB receptors, and in particular the CD8αβ-BBx2, strongly induced GM-CSF secretion. Notably, IL-13 secretion was observed in CD8+ cells expressing CD8-41BB receptors, which is characteristic of active 41BB signaling^30^. Similarly, CD4+ cells expressing CD8αβ-BBx1 or CD8αβ-BBx2 showed increased IFN-γ and GM-CSF secretion relative to CD8αβ. Intriguingly, CD4+ cells expressing CD8αβ had notably elevated IL-17A secretion, which was in marked contrast to the CD8αβ-BBx1 or CD8αβ-BBx2 groups. The cytokine profiles in the CD4+ cells suggest differential Th-like responses, which may be determined by the identity of the CD8α intracellular domain, where Lck interaction with the wild-type CD8α intracellular domain drives a Th1/Th17-like response, while CD8α-41BB fusions impart a stronger and more focused Th1 response.

### 3.4 CD8-41BB fusions promote regression of established tumors in challenging animal models

Given the marked *in vitro* functional enhancement conferred by the CD8αβ-BBx1 and CD8αβ-BBx2 receptors, we next performed *in vivo* studies using the previously described OVCAR3 xenograft model, in which the WT1-TCR alone cannot control established tumors (∼200 mm^3^) in NOG-IL15 mice **(Figure 4a)**. To further mitigate the immunosuppressive effects within the tumor microenvironment mediated by OVCAR3 expression of TGFβ1, T cells engineered for these studies also included CRISPR-Cas9-mediated knockout of TGFBR2. WT1-TCR T cells with TGFBR2 KO were resistant to the suppressive effects of TGFβ1 *in vitro* **(Figure S3a,b)**; however, TGFBR2 KO alone was not sufficient to slow tumor progression *in vivo* in this model **(Figure S3c)**. Consistent with the previous study, a dose of 1×10^7^ WT1-TCR T cells incorporating CD8αβ expression and TGFBR2 KO provided only partial and transient control of tumor progression **(Figure 4b)**. Strikingly, WT1-TCR/TGFBR2 KO cells with expression of either of the CD8αβ-BBx1 or CD8αβ-BBx2 receptors mediated complete and durable tumor regression in all treated mice, persisting throughout the duration of the study (110 days post tumor engraftment) **(Figure 4b,c)**. Given the robust response, we further evaluated T cell potency by treating the TCR KO control group when tumors reached ∼700mm^3^ with a single dose of WT1-TCR/CD8αβ-BBx1/TGFBR2 KO cells. Remarkably, all mice underwent a rapid and complete regression of these large tumors **(Figure 4d)**.

**Figure 4.**
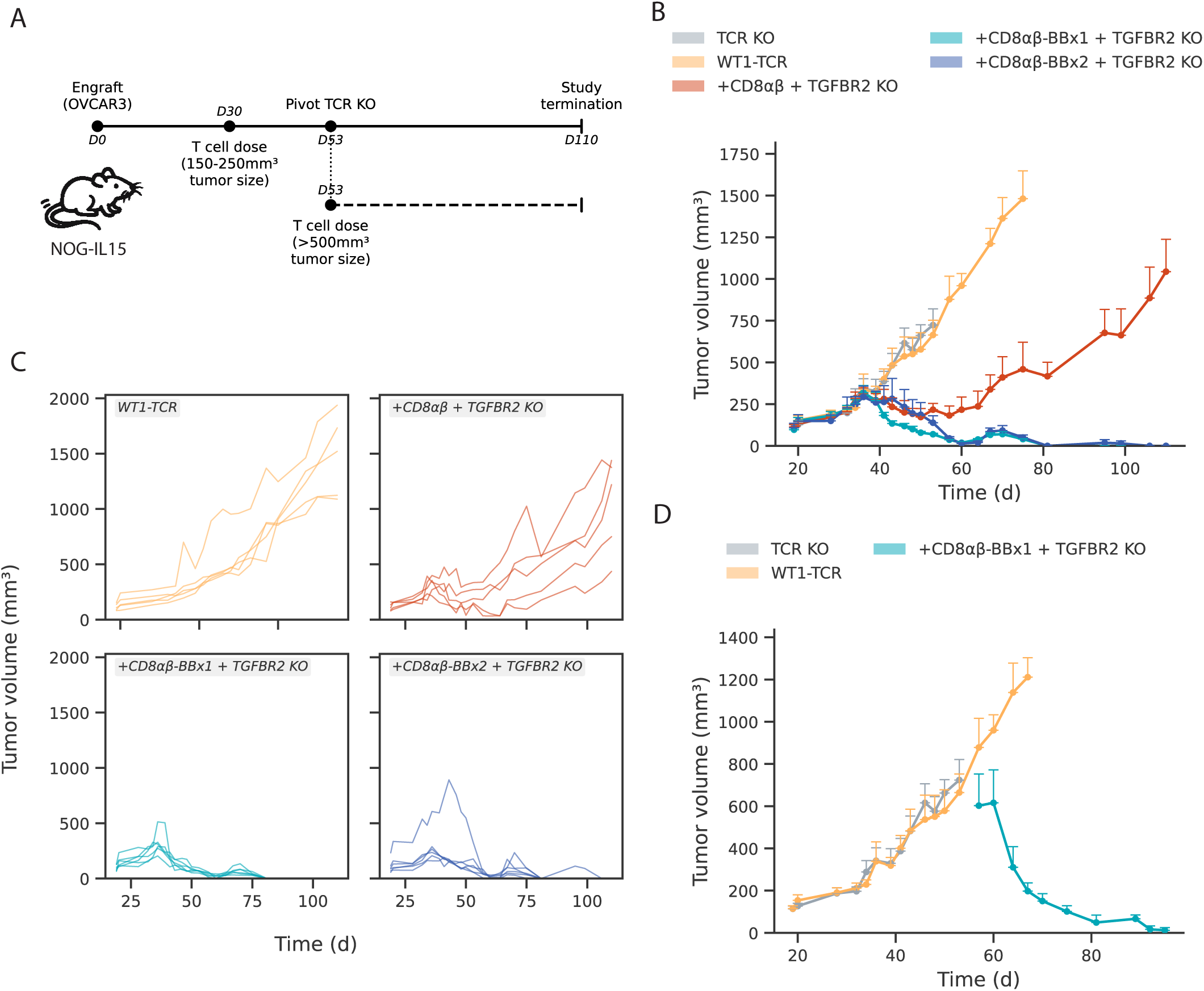
CD8-41BB fusions promote regression of established tumors in an ovarian cancer mouse model. **(a)** Schematic of OVCAR3 study design illustrating timing of subcutaneous (s.c.) tumor establishment and T cell dosing in NOG-IL15 mice. **(b)** Tumor volume measurements showing complete regression of OVCAR3 tumors following treatment with WT1-TCR/TGFBR2 KO T cells expressing CD8αβ-BBx1 or CD8αβ-BBx2 (n=5-6/group). **(c)** Individual tumor growth curves for mice in all treatment groups, showing tumor regression in all CD8αβ-BBx1 or CD8αβ-BBx2 treated mice with durable tumor control through study endpoint. **(d)** Fifty-three days post tumor engraftment, mice initially treated with TCR KO negative control T cells and bearing large tumors (mean tumor volume=723mm^3^) were treated with WT1-TCR/TGFBR2 KO T cells expressing CD8αβ-BBx1. Tumor volume measurements showed rapid tumor regression, with all mice achieving a complete response through study endpoint. Symbols in (b) and (d) represent mean ± SEM.

In the above studies, we used NOG-IL15 mice to provide human cytokine support for T cell engraftment and persistence. While in the clinical setting, lymphodepleting chemotherapy does induce an upregulation of IL-15 in serum, this effect is transient, waning over 1-2 weeks^31^. Since this contrasts with the constant IL-15 secretion in the NOG-IL15 mice, we proceeded to evaluate whether our CD8-41BB chimeras could mediate anti-tumor immunity in the absence of continuous IL-15 support. Additionally, in this study, we included treatment arms with high (7×10^6^ cells/mouse) and low (2×10^6^ cells/mouse) T cell dose levels to better compare efficacy between the CD8αβ-BBx1 and CD8αβ-BBx2 receptors **(Figure 5a)**. At the 7×10^6^ cells/mouse dose level, all mice treated with either CD8αβ-BBx1 or CD8αβ-BBx2 expressing WT1-TCR/TGFBR2 KO T cells had near complete tumor regression, confirming that continuous IL-15 support was not necessary **(Figure 5b)**. After complete tumor regression (∼50-days post T cell treatment), mice were rechallenged with another bolus of OVCAR3 cells at a distinct implantation site. Rechallenge tumors grew as small nodules, peaking at ∼100-150 mm^3^ over 10-15 days, before regression was observed in only the previously treated mice, suggesting the establishment of a durable memory immune response **(Figure 5c)**. In the 2×10^6^/mouse T cell dose level, we observed a significant improvement in tumor rejection with the CD8αβ-BBx1 chimera, with all CD8αβ-BBx1-treated mice achieving complete tumor regression, while the CD8αβ-BBx2-treated mice showed incomplete regression followed by slow outgrowth that continued throughout the study.

**Figure 5.**
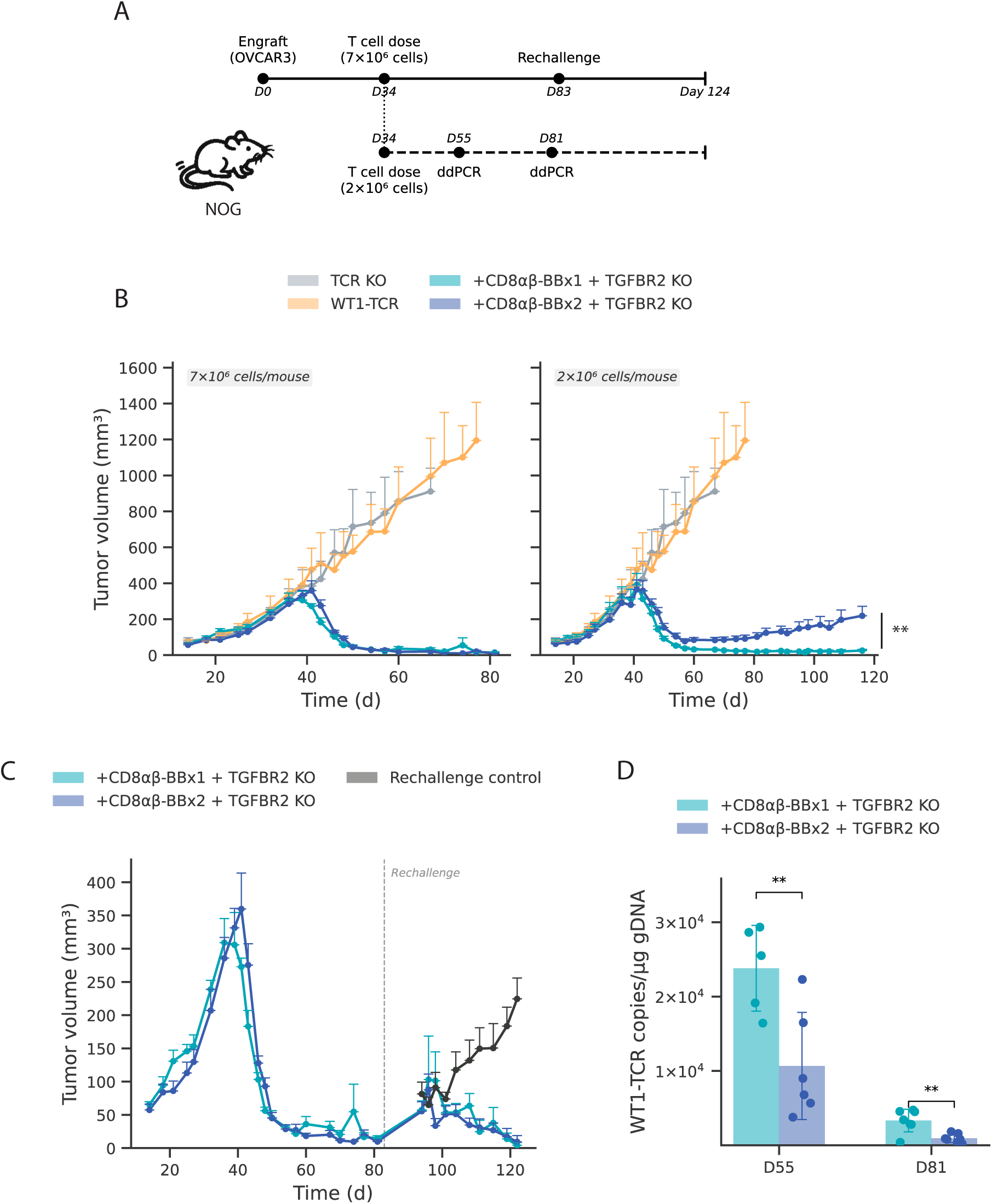
CD8αβ-BBx1 receptor expressing WT1-TCR cells have increased *in vivo* abundance and persistence to mediate long-term tumor control without IL-15 support. **(a)** Schematic of study design illustrating the establishment of OVCAR3 subcutaneous (s.c.) tumors in NOG mice followed by T cell dosing at high (7×10^6^ cells/mouse) or low (2×10^6^ cells/mouse) dose. Blood sampling for WT1-TCR pharmacokinetic (PK) analysis by droplet digital PCR (ddPCR) was performed in the 2×10^6^ cells/mouse cohort, while tumor rechallenge in the 7×10^6^ cells/mouse cohort at time points indicated in the schematic. **(b)** Tumor volume measurements showed complete tumor regression in both the CD8αβ-BBx1 and CD8αβ-BBx2 treatment groups at the high dose. In the low dose cohort, complete tumor control was observed only in the CD8αβ-BBx1 group. **(c)** Tumor volume measurements from mice in the 7×10^6^ cells/mouse cohort rechallenged with OVCAR3 cells on day 83showed continued tumor control. **(d)** WT1-TCR T cell abundance in mouse peripheral blood from the 2×10^6^ cells/mouse cohort was quantified by ddPCR, demonstrating significantly greater WT1-TCR T cell abundance in mice in the CD8αβ-BBx1 group relative to CD8αβ-BBx2. Each symbol represents an individual mouse (**p<0.01).

### 3.5 Improved T cell persistence and memory phenotype in CD8αβ-BBx1 expressing cells

The finding of improved *in vivo* tumor control with the CD8αβ-BBx1 co-receptor was surprising given that our *in vitro* data suggested superior activity with the CD8αβ-BBx2 receptor. To further evaluate this, we collected blood samples from the 2×10^6^ cells/mouse dose cohort at various points throughout the study to monitor T cell expansion and persistence using a droplet digital PCR (ddPCR) assay to quantify the WT1-TCR cell persistence. Notably, animals in the CD8αβ-BBx1 treatment group had significantly higher levels of circulating T cells at 21 days (study day 55) and 49 days (study day 81) post T cell treatment **(Figure 5d)** compared to the CD8αβ-BBx2 treatment group. Consistent with this finding, flow cytometry performed at study endpoint in the rechallenged cohorts identified a trend toward increased levels of WT1-TCR+, CD8+WT1-TCR+, and CD4+CD8+ WT1-TCR cells in the spleens of mice treated with CD8αβ-BBx1 receptor **(Figure S4a)**. Notably, a small (<5% of live cells) but distinctly defined population of T cells with a CD45RA+ CCR7+ CD28+ CD62L+ CD39-stem cell memory T cell (Tscm) phenotype was found in both the CD8αβ-BBx1 and -BBx2-treated groups **(Figure S4b)**, consistent with the finding that these receptors can mediate long-term durable tumor control in the rechallenge setting.

The increased *in vivo* abundance of CD8αβ-BBx1-expressing cells relative to CD8αβ-BBx2-expressing cells suggested reduced cell fitness in the latter population. To explore this, multiparameter flow cytometry was performed on the engineered cell product from three T-cell donors. Clustering using tSNE and FlowSOM algorithms **(Figure 6a)** showed a less favorable memory phenotype in the CD8αβ-BBx2-expressing cells. Specifically, in CD4+CD8+ cells, there was a decreased frequency of cells in T cell cluster with a favorable phenotypic profile (Pop.0: CD28+CD62L+) and an increase in more differentiated cell populations characterized by downregulation of memory markers CD28 and CD62L (Pop. 6) and expression of CD39 (Pop. 5), a marker associated with exhausted or terminally differentiated T cells^32^. **(Figure 6b,c)**. Similar trends were observed in CD8αβ-BBx2-expressing CD8+CD4-T cells, showing an increased frequency of cells in populations characterized by loss of CD27 and CD28 expression and upregulation of CD39 (Pop. 1, Pop 5) **(Figure 6d-f)**. Concomitantly, there was a non-significant trend towards decreased frequency of CD8αβ-BBx2 cells with an early memory phenotype (Pop. 8: CD45RA+CD27+CD28+CD62L+CD39-). In contrast to CD8αβ-BBx2, T cells expressing CD8αβ-BBx1 showed a similar phenotypic profile as T cells expressing CD8αβ. No cells in any group were found to express the activation/exhaustion markers PD-1, TIM-3, or LAG-3. Collectively, these findings demonstrate that the CD8αβ-BBx2 receptor, containing two 41BB ICDs, drives accelerated T cell differentiation during *ex vivo* cell expansion, which correlates with reduced *in vivo* persistence and tumor control.

**Figure 6.**
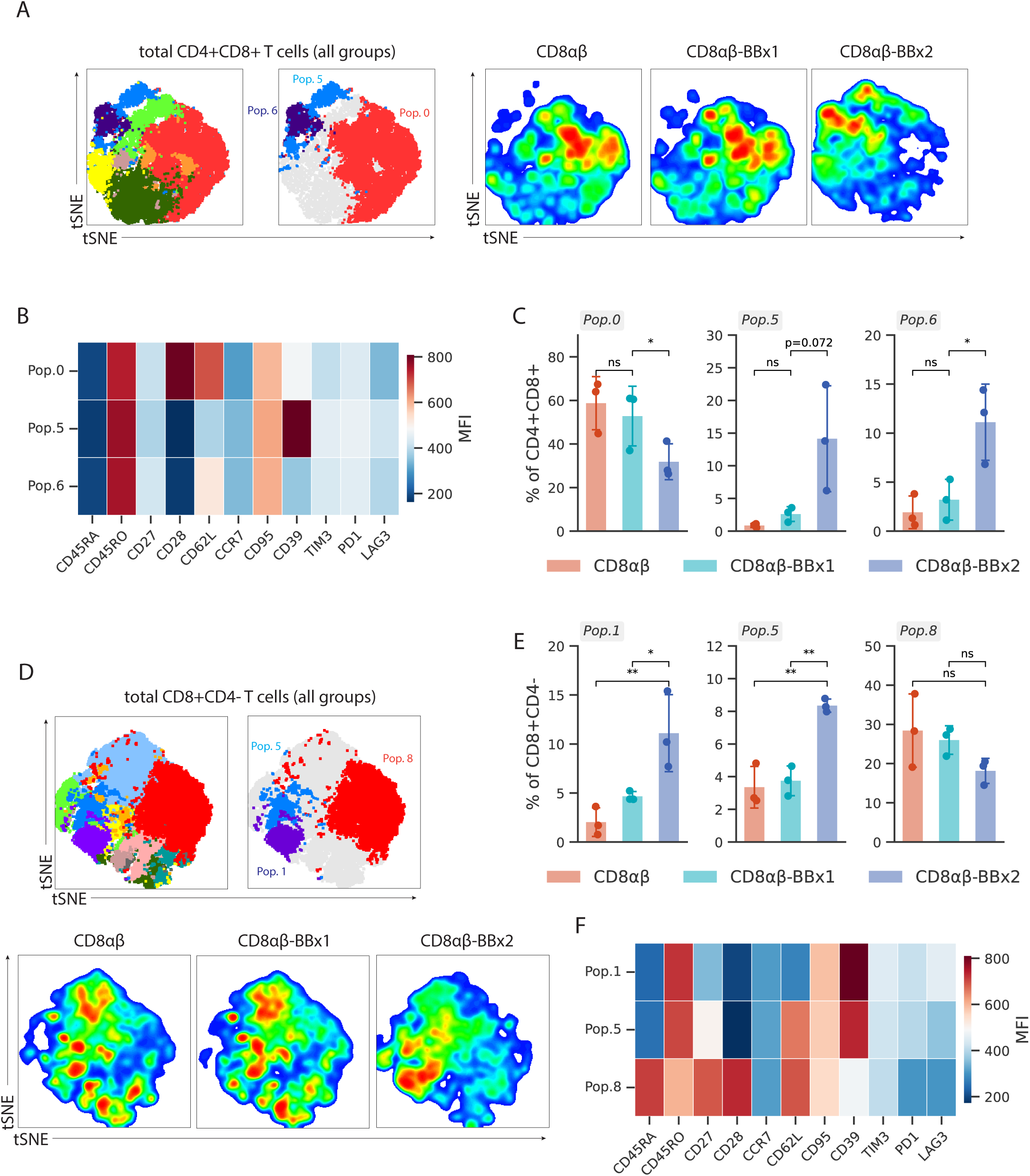
CD8αβ-BBx2 expression leads to a more differentiated memory phenotype. **(a)** The memory phenotype of engineered WT1-TCR T cells expressing the indicated CD8 co-receptor (n=3 donors) was characterized by multiparameter flow cytometry and unsupervised clustering using tSNE and FlowSOM algorithms. Cells were gated on live WT1-TCR+CD4+CD8+ T cells prior to clustering. **(b)** Heatmap showing median fluorescence intensity (MFI) of indicated markers across relevant populations (clusters identified in panel a). **(c)** Bar graphs depicting the frequency (%) of cells in each population across the three CD8 co-receptor groups. Each dot represents an individual donor (n=3, *p<0.05). **(d-f)** A similar analysis was performed after gating on CD8+CD4-cells with flow plots (d) showing cell clustering, bar graphs (e) showing frequency of cells in each population (n=3, *p<0.05, **p<0.01) and the heatmap (f) showing MFI of each phenotypic marker across the populations of interest.

### 3.6 CD8αβ-BBx1 enhances WT1-TCR T cell in vivo efficacy in an AML model

To test the WT1-TCR/CD8αβ-BBx1/TGFBR2 KO T cells against AML *in vivo*, we developed a disseminated tumor model using the OCI-M1 cell line. Here, luciferized OCI-M1 cells were injected intravenously (i.v.) into previously irradiated NOG mice and tumor engraftment and distribution were monitored by IVIS bioluminescence imaging. Strikingly, when mice were treated 10 days after tumor establishment (luciferase signal ∼2×10^7^ photons/second (p/s)) with WT1-TCR/CD8αβ-BBx1/TGFBR2 KO T cells, complete tumor elimination was observed in all mice **(Figure 7a,b)**. In contrast, WT1-TCR/TGFBR2 KO T cells expressing wild-type CD8αβ conferred only transient tumor growth inhibition, comparable to that observed in WT1-TCR/TGFBR2 KO T cells without co-receptor expression. This data further confirms a potency enhancement conferred by the 41BB signaling domain in both solid and hematological tumor models.

**Figure 7.**
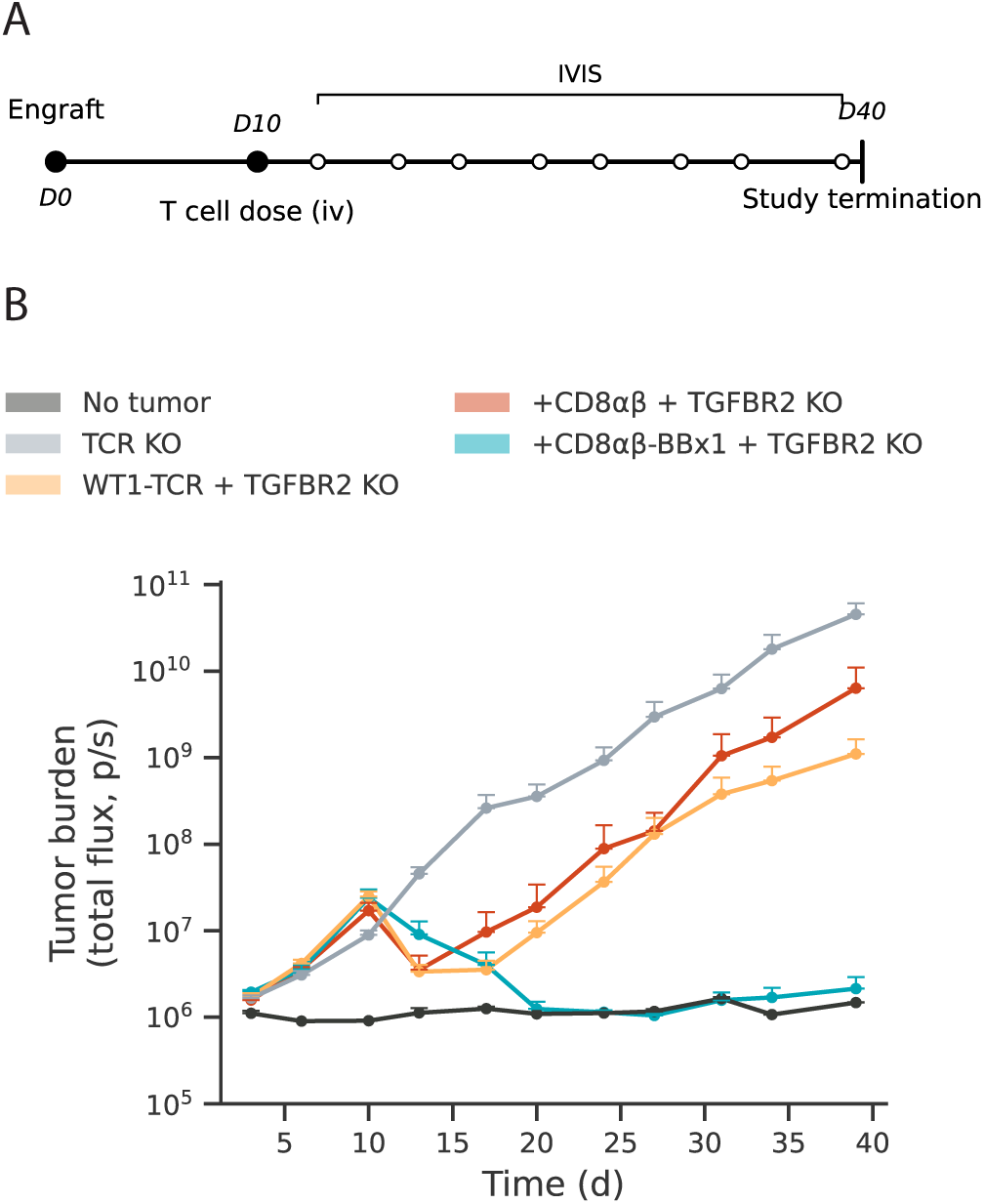
WT1-TCR T cells expressing CD8αβ-BBx1 receptor regressed established tumors in a disseminated AML mouse model. **(a)** NOG mice were irradiated prior to intravenous (i.v.) injection of OCI-M1-GFP/luciferase (GFP/luc) cells. Ten days post tumor engraftment, animals were randomized and treated with the indicated T cell group. IVIS bioluminescence imaging was used to monitor tumor burden over time; representative images acquired at indicated time points are shown. **(b)** Quantification of luciferase signal showed complete regression of OCI-M1 tumors following treatment with WT1-TCR/TGFBR2 KO T cells expressing CD8αβ-BBx1 (n=6-8/group); arrow indicates the time of T cell dosing.

### 3.7 WT1-TCR specificity and HLA-A*02:01 restriction is consistent upon CD8-41BB mediated enhancement

Next, we sought to assess whether the enhancement of WT1-TCR T cells conferred by CD8-41BB co-expression came at a cost of reduced target specificity or loss of HLA-A*02:01 restriction. First, we confirmed that CD8 or CD8-41BB-expressing WT1-TCR T cells showed no reactivity toward HLA-null K562 cells, which endogenously express WT1 **(Figure S5a)**. A similar approach was taken to confirm WT1-dependent killing using a WT1-negative, HLA-A*02:01+ H441 lung adenocarcinoma cell line. As expected, these cells were refractory to killing by all T cell groups unless transduced to express WT1 **(Figure S5b)**. We next evaluated the ability of enhanced WT1-TCR/CD8-41BB T cells to mediate killing of primary AML cells with and without HLA-A*02:01 expression. Efficient tumor cell lysis was observed across the two HLA-A*02:01+ primary AML samples tested, with no lysis of HLA-A*02:01^−^ cells, further confirming HLA-restricted activity **(Figure S5c)**. Notably at the low E:T ratio tested here (1:12) one of the HLA-A*02:01+ pAML samples was refractory to killing with WT1-TCR T cells but was killed effectively with WT1-TCR T cells when armored with either CD8-41BB receptor and TGFBR2 KO.

Additionally, we sought to evaluate whether the CD8 costimulatory expression altered specificity toward the targeted WT1 peptide epitope. Our previous study used peptide pulsing of alanine mutated WT1_37–45_ peptides to define the key recognition motif for this WT1-TCR as VLD(F)APxxA, where A is a natural alanine that was not investigated, x is not critical for binding, and position F4 induced T cell reactivity only when pulsed at supraphysiological concentrations (500nM)^20^. To assess any changes in specificity upon CD8-41BB co-receptor expression, H441-GFP cells were pulsed with the same panel of peptides. Encouragingly, when alanine-substituted peptides were pulsed at a concentration representing the EC90 value for the WT1_37–45_ peptide, co-receptor expressing WT1-TCR T cells retained the same VLD(F)APxxA specificity profile as previously described **(Figure S5d)**. Lastly, we sought to assess the on-target, off-tumor reactivity of CD8- or CD8-41BB-enhanced WT1-TCR T cells against normal cells. WT1 is expressed by several healthy tissues, including CD34^+^ hematopoietic cells, at levels 100 to 10000-fold lower than that observed in AML or ovarian cancer^33^. Our previous data showed that the WT1-TCR alone did not trigger T cell responses in CD34^+^ HSC co-cultures, but the increased avidity and potency conferred by overexpression of CD8 or CD8-41BB may heighten antigen sensitivity. Reassuringly, when CD8-41BB co-expressing WT1-TCR T cells were co-cultured with HLA-A*02:01^+^ and HLA-A*02:01^−^ CD34^+^ cells, no significant cytokine secretion was detected in any group **(Figure S5e)**. We also confirmed that the CD8-41BB co-expressing WT1-TCR/TGFBR2 KO T cells did not show cytotoxicity against HLA-A*02:01+ healthy donor bone marrow mononuclear cells (BMMCs) **(Figure S5f)**. These data suggest that the potency conferred by CD8 or CD8-41BB overexpression does not significantly impact the intrinsic specificity or HLA restriction determined by the TCR.

### 3.8 CRISPR-based targeted insertion of CD8-41BB and TCR chain into the *TRAC* locus from a single donor template

While efficient, the dual targeted knock-in (TKI) process used to engineer T cells with the TCR and CD8-41BB transgenes is complex, requiring delivery of two AAV-based HDR templates and an additional CRISPR-Cas9 editing step. Additionally, the dual TKI strategy introduces insertion variations, where some cells have integration of only one of the two transgenes. Thus, we explored whether the HDR template design could be modified to allow for a single-step insertion of both a CD8αβ-41BB co-receptor and the WT1-TCR into the *TRAC* locus, while accommodating the upper size limit of AAV6 (∼4.7 kb). To accomplish this, the CD8αβ-BBx2 and TCR transgenes were cloned into a single backbone (separated by an additional 2A self-cleaving peptide site), and the transgenic TCRα chain within the HDR cassette was designed such that, upon in-frame insertion, the endogenous *TRAC* 3’ coding sequence and UTR would complete TCR expression^34^. Encouragingly, CRISPR-Cas9-mediated single TKI using AAV6 transduction of the modified template was highly efficient, with >80% of T cells expressing the transgenes **(Figure 8a)**. As expected, the single TKI approach yielded a high proportion of cells with homogeneous expression of both transgenes, whereas the dual TKI approach produced a mixture of cells expressing either one or both transgenes **(Figure 8b)**.

**Figure 8.**
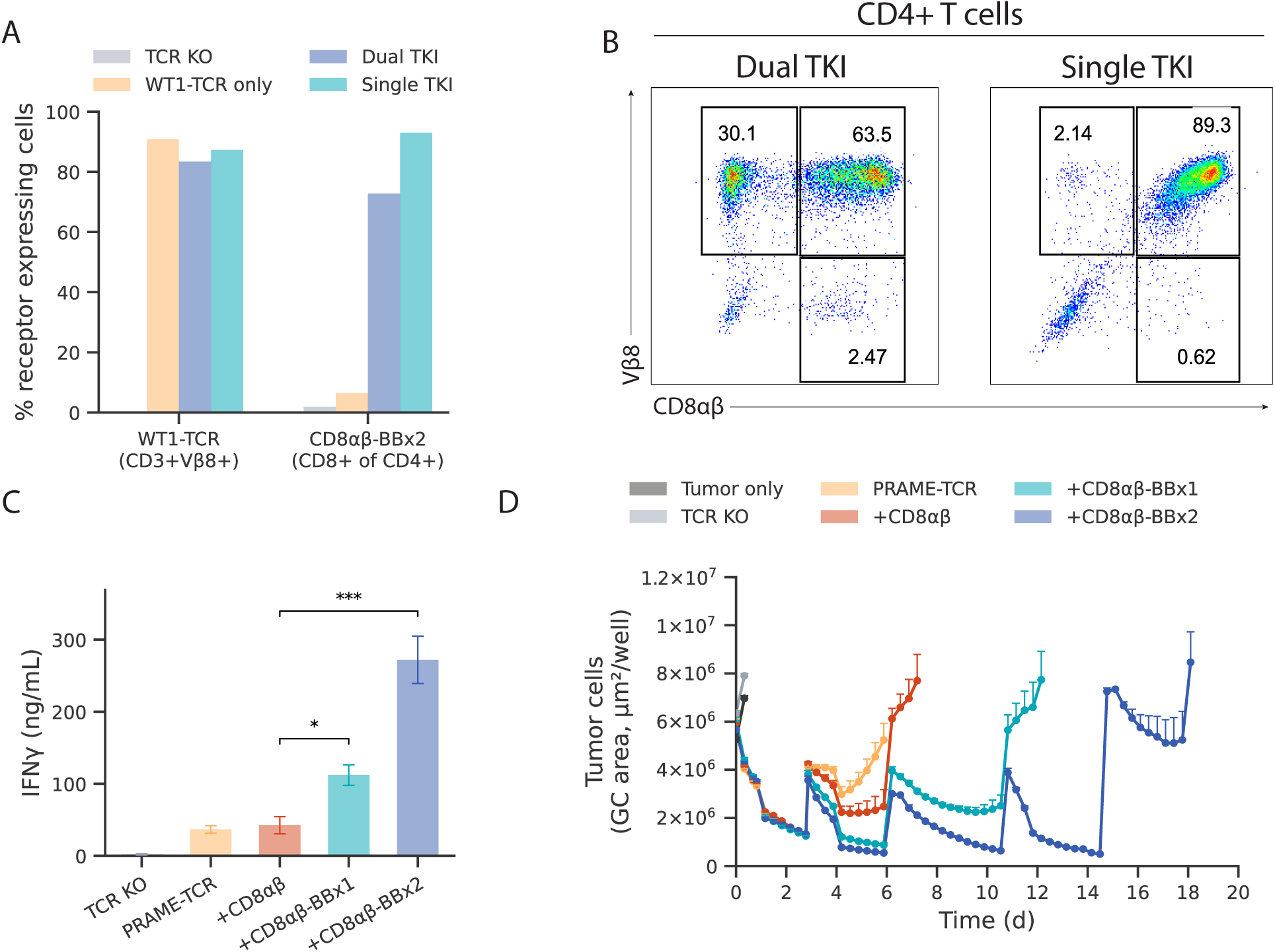
Single step targeted integration of both CD8-41BB co-receptor and TCR into *TRAC* locus and application to a second TCR. **(a)** Single targeted knock-in (TKI) using a redesigned AAV template encoding both the WT1-TCR and CD8αβ-BBx2 transgenes was compared to the dual TKI approach. Bar graphs showed the percentage of CD4+ T cells expressing the WT1-TCR (CD3+Vβ8+) or CD8αβ-BBx2. **(b)** Representative flow cytometric plots demonstrating the correlation between WT1-TCR and CD8αβ-BBx2 expression in the dual vs single TKI approaches. **(c)** T cells engineered with a PRAME-directed TCR and the indicated CD8 co-receptors were assayed for IFN-γ release and **(d)** tumor cytotoxicity in a serial rechallenge assay with GFP-labelled A375 cells. The CD8αβ-BBx1 and CD8αβ-BBx2 co-receptors conferred functional enhancement with the PRAME-TCR, consistent with the observations for WT1-TCR.

### 3.9 CD8-41BB fusions enhance the activity of a second high affinity TCR

Lastly, we sought to evaluate the applicability of the CD8-41BB co-receptors as a general enhancement strategy for other clinically relevant TCRs and targets beyond WT1. To this end, we generated a single HDR template incorporating an HLA-A*02:01-restricted TCR targeting PRAME, a target currently being investigated in solid tumor clinical trials^35,36^. PRAME-TCR-T cells expressing CD8 receptors were efficiently engineered and challenged *in vitro* with the A-375 melanoma cell line (PRAME+, HLA-A*02:01+). Consistent with the results obtained using the WT1-TCR, PRAME-TCR-T cells expressing CD8αβ-BBx1 or CD8αβ-BBx2 co-receptors had significantly increased IFN-γ secretion and enhanced tumor cytotoxicity in serial rechallenge assays **(Figure 8c,d)**. The consistent functional enhancement observed across both WT1- and PRAME-targeting TCRs supports the broader utility of CD8-41BB co-receptor expression as a universal strategy for enhancing HLA class I-restricted TCR-T cell therapies.

## 4 Discussion

The advent of second-generation CARs has led to remarkable clinical breakthroughs, providing life-changing treatment options for patients with certain hematological cancers, and has invigorated a field of research to apply cell therapies across other cancer types and, more recently, autoimmune diseases. Key to these receptors was the incorporation of CD28 or 41BB intracellular domains to transduce co-stimulatory signals in an antigen-dependent manner^37–39^. Additionally, it is now well established that CD4+ CAR-T cells play a critical role in shaping anti-tumor responses by mobilizing endogenous immune cell subsets and supporting transferred CD8+ CAR-T cells^40^. Utilizing CRISPR-Cas9 and synthetic biology, we sought to bring these valuable properties to TCR-T cells, which, by their nature, have unique advantages in terms of broader antigen accessibility and increased antigen sensitivity.

To accomplish this, we screened several CD8-41BB and CD8-CD28 chimera designs. Given our initial experiments demonstrated poor activity from CD8αα homodimers, subsequent construct designs focused on CD8αβ heterodimers. This screening approach led to the identification of two CD8-41BB receptors that imparted substantially increased activity to T cells co-expressing WT1- or PRAME-targeting TCRs. Notably, both active constructs identified in this screen have 41BB replacement of the CD8α intracellular domain. This data contrasts with the recent report of a CD8β-CD28 fusion which demonstrated improved *in vitro* and *in vivo* anti-tumor functions in the context of a MAGE-A1 specific TCR^41^. However, this study did not report evaluation of a CD8α-41BB chimera, based on early findings of significantly reduced expression of CD8α-CD28 constructs compared to CD8β-CD28 in Jurkat cells. Notably, we did not observe significant expression differences across CD8α or CD8β fusion proteins upon ICD replacement with CD28 or 41BB, which may be explained by the CRISPR TKI method used here rather than lentiviral vector delivery of the combined CD8αβ and TCRαβ chains. Other explanations could include use of different promoters (EF1α vs MSCV), codon optimization variations, and primary T cells vs immortalized Jurkat cells. While these differences in cell engineering strategies may explain why CD8α-modifications enabled high expression and function in this study, they do not currently explain why the CD8β-CD28 fusion tested here did not demonstrate the enhanced function over wild-type CD8αβ as reported. Future studies are needed to mechanistically characterize and expand the design space of CD8 fusion receptors.

In CD4+ T cells expressing the CD8 co-receptors, replacement of the CD8α-Lck ICD with the 41BB ICD led to a differentiated cytokine profile, most notably characterized by reduced IL-17A secretion and increased IFN-γ secretion. High levels of IL-17A secretion from CD4+ T cells expressing CD8αβ has not been previously reported; however, our data suggest that this is likely driven by CD8α-Lck signaling, which is attenuated upon substitution of the CD8α ICD with the 41BB ICD. In CD8+ T cells, addition of CD8α-41BB led to increased cell surface expression of CD8αβ heterodimers and significantly enhanced antigen-dependent T cell proliferation, cytokine secretion, and sustained tumor cell killing. In contrast, additional expression of wild-type CD8αβ provided only a marginal benefit, which may be attributed to increased CD8 surface expression enhancing TCR-pMHC binding avidity. Notably, the CD8+ T cells expressing CD8α-41BB would simultaneously have the capacity for both endogenous CD8-Lck and transgenic CD8-41BB signaling. Further studies, including knockout of the endogenous CD8α chain, are required to delineate the relative impact of each signaling pathway; nonetheless, our data point to a clear benefit conferred by the antigen driven 41BB co-stimulation in both CD4+ and CD8+ T cells.

The *in vitro* assays testing the CD8αβ-BBx1 and CD8αβ-BBx2 chimeras suggested increased tumor cell reactivity and killing with the CD8αβ-BBx2 chimera. While both constructs showed robust *in vivo* tumor control, it was initially surprising to observe superior *in vivo* efficacy with CD8αβ-BBx1 when tested at a low T cell dose. This superior efficacy was correlated with increased WT1-TCR T cell abundance in the blood and spleens of these mice. This discrepancy between the *in vitro* and *in vivo* experiments may be explained in part by the CD8αβ-BBx2 receptor driving a more differentiated T cell phenotype in CD4+CD8+ and CD8+ T cells **(Figure 6)**.

Cumulatively, these studies highlight the benefit of engineering TCR-based T cell therapies with CD4+ T cell helper activity and built-in co-stimulatory signaling. Other approaches to impart T cells with co-stimulation include chimeric co-stimulatory receptors that signal upon binding tumor antigens (e.g. FOLR1^42^, CD19^43^), switch-receptors that drive positive signals upon engaging inhibitory ligands (e.g. CD200^14^, PD-1^12,13^, FasL^15^), and a more recently described CD80-41BB fusion receptor that can be agonized by either CD28 or CTLA-4^44^. While preclinical studies have demonstrated the potency benefits conferred by co-stimulation through these chimeric receptors, the use of cell line models expressing homogeneous levels of the respective counter-ligand may not reflect the heterogeneous expression patterns observed in human cancers. In contrast to these approaches, armoring TCR-T cells with CD8-41BB receptors leads to co-stimulatory signal transduction in an antigen-dependent manner, as is the case with 2^nd^-generation CARs, while also having the benefit of promoting CD4+ T cell activity.

## 5 Conclusions

In summary, our data support the use of CD8-41BB receptors to enhance the potency of TCR-T cells, which could represent a potential next-generation approach for developing potent cell therapies for tumor targets that are inaccessible to CAR-T cell therapies. Notably, the engineering of T cells through expression of transgenic TCRs and CD8 fusion receptors, combined with targeted gene knockout to prevent TME-mediated immunosuppression (e.g. TGFBR2 KO), was readily achieved utilizing CRISPR-Cas9-based genome editing, providing a promising platform for future clinical development. Moreover, the benefit of this approach in enhancing not only CD4+ T cells, but also CD8+ CTLs supports further investigation of the safety and potency of CD8-41BB receptors for other TCR-based anti-tumor modalities, such as *ex vivo*-expanded tumor-infiltrating lymphocytes (TILs) or circulating tumor-reactive lymphocytes^45,46^.

## 6 Materials and Methods

### 6.1 Cell Lines

ALL cell line 697 (ACC 42), AML cell lines OCI-AML3 (ACC 582) and OCI-M1 (ACC 529) were obtained from Leibniz Institute-German Collection of Microorganisms and Cell Cultures GmbH (DSMZ). The lung adenocarcinoma cell line NCI-H441 [H441] (HTB-174), melanoma cell line A-375 [A375] (CRL-1619), and ovarian adenocarcinoma cell line NIH:OVCAR-3 [OVCAR3] (HTB-161) were obtained from the American Type Culture Collection (ATCC). K562-Fluc-Puro (Cat# CL171) and K562-HLA-A2-Luc2/Puro (Custom order) cell lines were obtained from Imanis Life sciences. OCI-M1, OCI-AML3, and A375 were cultured in DMEM media (Thermo Fisher Scientific) supplemented with 20% FBS (OCI-M1) or 10% FBS (OCI-AMl3, A375) (Thermo Fisher Scientific). K562-Fluc-Puro, K562-HLA-A2-Luc2/Puro, and H441 were cultured in RPMI-1640 (Thermo Fisher Scientific) supplemented with 10% FBS. OVCAR3 was cultured in RPMI-1640 supplemented with 20% FBS and 0.01 mg/mL bovine insulin (Sigma-Aldrich; Cat# I0516). All cell lines were maintained and passaged according to the vendor’s recommendations. To generate cell lines stably expressing GFP and luciferase, cells were transduced with either a GFP-encoding lentivirus or a reporter lentivirus encoding both GFP and luciferase (LV-SFFV-Luc2-P2A-EmGFP; Imanis Life Sciences, Cat# LV050L) and subsequently sorted by GFP expression. The H441 cell line was transduced with a WT1-GFP-encoding lentivirus and sorted by GFP expression for the generation of the H441-WT1-GFP cell line. For *in vivo* experiments, OVCAR3 was transduced with lentivirus (LV-Luc2-P2A-Puro; Imanis LV012L) and selected by serial passaging in puromycin-containing media.

### 6.2 Animals

Female NOG (NOD.Cg-Prkdcscid Il2rgtm1Sug/JicTac) and NOG-IL15 (NOD.Cg-Prkdcscid Il2rgtm1Sug Tg(CMV-IL2/IL15)1-1Jic/JicTac) mice were obtained from Taconic Biosciences (Albany, NY, USA) and were aged ≥6 weeks at the time of experimentation.

### 6.3 Design of HDRT Constructs

The design of the homology-directed repair template (HDRT) and associated AAV6 production for targeted integration of the WT1-TCR in the *TRAC* locus have been described previously^20^. HDRTs enabling insertion of wild-type or chimeric CD8 chains in the *AAVS1* locus were designed with 500 bp homology arms (complementary to the *AAVS1* sgRNA target site) flanking the genes of interest (GOIs) driven by the EF1α promoter. The CD8α and CD8β chains were separated by a self-cleaving 2A peptide sequence. CD8α, CD8β, 41BB (CD137), and CD28 sequences, including annotation of extracellular, transmembrane, and intracellular domains used for construct designs, were sourced from UniProt (P01732, P10966, Q07011, P10747). Single AAV HDRTs encoding both TCR and CD8 transgenes were designed similarly to the original WT1-TCR HDRT by appending the respective CD8 chimeras to the 5’ end of WT1- or PRAME-TCR coding sequences, separated by additional self-cleaving 2A peptide sequences. However, to accommodate AAV6 packaging size limits, the EF1α promoter was replaced with a smaller EF1α-core promoter, and the 3’ coding sequence and UTR of the endogenous *TRAC* gene were utilized to complete expression of a the TCRα chain upon in-frame insertion of the transgene. All gene synthesis and cloning was performed by GenScript (New Jersey, USA) and all constructs were sequence-verified prior to use. Methods for AAV6 production has been described elsewhere^20^.

### 6.4 T Cell Isolation

Healthy donor leukopaks were obtained from Cell Solutions, Charles River Laboratories. CD4+ and CD8+ T cells were isolated by positive selection using the MultiMACS™ Cell24 Separator Plus (Miltenyi Biotec; Cat# 130-098-637). Briefly, each leukopak was resuspended in separation buffer (PBS, 2% FBS, 1 mM EDTA) and incubated with CD8+ microbeads (Miltenyi Biotec; Cat# 130-045-201). After incubation, the labeled CD8+ fraction was captured by the magnetic Multi-24 Column Blocks (Miltenyi Biotec; Cat# 130-095-691) and recovered. The initial flow through was retained and labelled with CD4+ microbeads and CD4+ T cells were isolated using the same magnetic separation procedure described above. The obtained CD8+ and CD4+ T cells were cryopreserved using CryoStor® CS10 (StemCell Technologies; Cat# 07930).

### 6.5 CRISPR-Cas9 based T Cell Engineering

Previously isolated CD4+ and CD8+ T cells were thawed and resuspended in a 1:1 ratio in T cell growth media (TCGM) consisting of CTS™ OpTmizer™ T Cell Medium (Thermo Fisher Scientific) supplemented with 2.5% human serum (BioIVT), 1% GlutaMax (Thermo Fisher Scientific), 1% HEPES (Thermo Fisher Scientific), 1% Penicillin-Streptomycin (Thermo Fisher Scientific), 10 ng/mL IL-2 (PeproTech), 5 ng/mL IL-7 (PeproTech), and 5 ng/mL IL-15 (PeproTech)). After overnight rest, T cells were activated using TransAct (Miltenyi Biotec, Cat# 130-111-160). Unless otherwise specified, CRISPR-Cas9 gene knockout was achieved as previously described^48^ by transfection of activated T cells with lipid nanoparticles (LNPs) containing Streptococcus pyogenes Cas9 (Sp. Cas9) mRNA and sgRNA targeting *TRBC1/2* or *TGFBR2* as indicated. LNP delivery of genome editing components has been demonstrated to be an efficient and non-toxic approach for *ex vivo* gene editing^49^. For targeted gene insertion, T cells were transfected with LNPs containing Sp. Cas9 mRNA and sgRNA targeting *TRAC* or *AAVS1* simultaneously with AAV6 transduction of HDRTs in the presence of a small molecule DNA-PK inhibitor. 4-5 days post activation, T cells were transferred to GREX24 (Wilson Wolf Manufacturing; Cat# 80192M) or GREX6 (Wilson Wolf Manufacturing; Cat# 80240M) well plates and expanded according to the manufacturer’s recommendations. Cells were harvested 10-11 days after activation and cryopreserved in CryoStor^®^ CS10 for future use.

### 6.6 Serial Rechallenge Assays

Engineered T cells were thawed and rested overnight in TCGM prior to co-culture with GFP-labelled tumor cells (OVCAR3, OCI-M1, or A375; 1×10^5^ cells/well) at a 4:1 E:T ratio (unless otherwise noted) in cytokine-free TCGM. For subsequent tumor challenges, media was carefully removed from the top of each well every 72-96 hours for addition of fresh tumor cells (1×10^5^ cells in 100µL/well). Tumor cell lysis was monitored continuously using the Incucyte^®^ S3 Live-Cell Analysis Systems (Sartorius), whereby a reduction of GFP fluorescence intensity or GFP-positive area was used to quantify tumor cell lysis.

### 6.7 Cytokine Analysis

Cell culture supernatants were collected 24h post tumor challenge unless otherwise specified. Cytokine secretion was quantified using Mesoscale Discovery (MSD) U-PLEX CAR-T Cell Combo 1 (hu) (Cat# K15338K-2) or U-PLEX Custom Biomarker (hu) Assays, containing GM-CSF, IFN-γ, IL-2, IL-4, IL-10, IL-13, IL-17A and TNF-α (Cat# K15067M-2) according to the manufacturer’s recommendations, and data were analyzed using the Discovery Workbench software.

For the bone marrow toxicity study, samples from three cryopreserved, positive-selected CD34+ bone marrow cell donors were obtained (AllCells). WT1-TCR T cells were co-cultured with CD34+ bone marrow cells or OVCAR3 tumor cells at an E:T ratio of 1:2.5 for 24h, after which supernatants were collected. IFN-γ levels were quantified using the Mesoscale Discovery (MSD) U-PLEX IFN-γ Assay (hu) (Cat# K151TTK-2) according to the manufacturer’s recommendations.

### 6.8 CD4+CD8+ and CD8+ cell enrichment

For independently assessing the functionality of engineered CD4+, CD4+CD8+, and CD8+ T cells, we separated the fractions using CD4+ microbeads (Miltenyi Biotec; 130-045-101). Briefly, overnight rested T cells were washed and resuspended in separation buffer, incubated with CD4+ microbeads for 15 mins at 4°C. Cells were then washed and passed through a magnetic LS column (Miltenyi Biotec, Cat# 130-042-401) mounted on the QuadroMACS separator (Miltenyi Biotec, Cat# 130-090-976). The flow-through (CD8+ T cell fraction) and eluate (CD4+ T cell fraction) were collected, and the cell purity of each fraction was confirmed by flow cytometry.

### 6.9 Flow cytometry

A list of antibodies used in this study is provided in **Table S1**. For surface staining, T cells were resuspended in FACS buffer (PBS, 2% FBS, EDTA 2 mM) containing diluted antibodies and stained at room temperature for 30 mins. Post staining, T cells were washed, resuspended in FACS buffer, and subsequently processed on a CytoFLEX flow cytometer (Beckman Coulter). For WT1-TCR detection, either an anti-Vβ8-PE antibody (BioLegend) or VLDFAPPGA peptide-MHC Dextramer-PE reagent (Immudex) was used. The CD8αβ heterodimer on CD4+ T cells post CD8 transgene insertion was quantified using antibody clones 2ST8.5H7(BD Biosciences) or QA20A40 (BioLegend). Mean fluorophore intensity (MFI) of CD8 in the transgene-in serted CD4+CD8+ population was used to quantify CD8 surface expression levels.

For intracellular cytokine analysis, engineered T cells were co-cultured with tumor cells in a 96-well U-bottom in medium containing manufacturer recommended concentrations of GolgiPlug (BD Biosciences; Cat# 555029) and GolgiStop (BD Biosciences; Cat# 554724). Following 16 hours of co-culture or cell activation, T cells were stained with a fixable viability dye, followed by incubation with a cocktail of surface antibodies targeting CD3, CD4, CD8, and Vβ8. Cells were then fixed and permeabilized according to the manufacturer’s protocol (Fixation Medium (Medium A), Cat# GAS001S100; Permeabilization Medium (Medium B), Cat# GAS002S100; Thermo Fisher Scientific) and then intracellular cytokines were labelled using anti-IFN-γ and anti-TNF-α antibodies.

For flow-based cytotoxicity assays, CellTrace™ Violet (CTV, Invitrogen; Cat# C34557)-labelled engineered T cells were co-cultured at an E:T ratio of 1:12 with primary AML blasts for 4 days. Post co-culture, live AML blasts were quantified by flow cytometry by enumerating CTV-HLA-A2+ events (HLA-A*02:01+ AML blasts) or CTV-CD33+ (HLA-A*02:01-AML blasts) events to assess the cytotoxicity of the engineered T cells. In a separate experiment, CTV-labelled engineered T cells were co-cultured at an E:T ratio of 1:12 with HLA-A*02:01+ primary AML blasts or HLA-A*02:01+ healthy donor bone marrow mononuclear cells (BMMCs; DLS) for 4 days. Post co-culture, live cells were quantified by flow cytometry by enumerating CTV-HLA-A2+ events, as described above.

All data analysis, including tSNE and FlowSOM clustering, was performed using FlowJo software package (v.10.6.1 or v.10.7.1) and associated analytical plugins.

### 6.10 Cytotoxicity Assays

Engineered T cells were thawed and rested overnight in TCGM. The following day, co-cultures with tumor cell lines were established at a range of E:T ratios. For luciferase-based cytotoxicity assays, after 48h of co-culture, residual luciferase activity, measured as relative light units (RLU), was quantified using the Bright-GLO Luciferase Assay System (Promega, Cat# E2610) and a CLARIOstar Plus plate reader (BMG Labtech). Percent tumor cell lysis was calculated using the following formula:

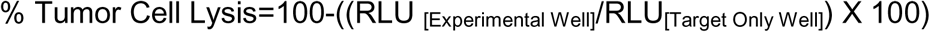

To assess the real-time cytotoxicity kinetics, the Incucyte^®^ S3 Live-Cell Analysis System (Sartorius) was used, whereby loss of GFP fluorescence from GFP-labelled tumor cell lines was used to quantify target cell lysis. GFP measurements were normalized to the tumor cell only control (100%) for each timepoint.

For the alanine scanning specificity study, the WT1-negative H441-GFP/Luc2-expressing cell line was pulsed with an 8-point dose titration of wild type VLDFAPPGA or individual 9-mer peptides in which each non-alanine residue was substituted with an alanine The WT1 peptide RMFPNAPYL was included as a negative control. WT1-TCR T cells were co-cultured with the peptide-pulsed H441-GFP/Luc2 cells at a 1:1 E:T ratio. Tumor cell lysis was monitored using the Incucyte® S3 Live-Cell Analysis Systems as described above. GFP measurements were normalized to the tumor cell only control (100%), and tumor cell lysis was reported at the 72h time point.

### 6.11 Proliferation Assays

Engineered T cells were labelled with CellTrace™ violet dye (CTV; Invitrogen; Cat# C34557) according to the manufacturer’s protocol and added to tumor cells in a 96-well flat-bottom plate at an E:T ratio of 2:1 in cytokine-free TCGM. After 4□days of co-culture, T cells were phenotyped by flow cytometry to assess proliferation. Briefly, T cells were stained with a cocktail of antibodies targeting CD3, CD4, CD8, and Vβ8. Viakrome 808 (Beckman Coulter; Cat# C36628) was included as a fixable viability dye. Stained T cells were resuspended in 90 μL FACS buffer and 10□μL CountBright™ Absolute Counting Beads (Invitrogen; Cat# C36995) were added to each well prior to analysis on a CytoFLEX flow cytometer (Beckman Coulter). T cells were gated based on size (FSC/SSC), T cell subset (CD4+CD8+ or CD8+CD4⁻), and WT1-TCR expression (Vβ8+ CD3+). Proliferating cells were identified by gating on CTVlow events, and the absolute number of proliferating cells was calculated using the following formula:

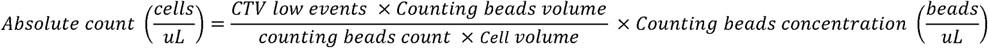

### 6.12 TGF-β1 Signaling and Suppression Assays

To assess the impact of TGFBR2 KO on TGF-β1-mediated SMAD signaling, engineered T cells or unedited controls were incubated with recombinant TGF-β1 (10 ng/mL; Sigma-Aldrich; Cat# 11412272001) for 30 mins to induce SMAD phosphorylation. Phosphorylated SMAD2/3 (pSMAD2/3) was assayed by intracellular flow cytometry. Briefly, cells were washed and resuspended in pre-warmed 1X BD Phosflow™ Lyse/Fix Buffer (BD Biosciences; Cat# 558049) and incubated for 10 mins at 37°C. Post incubation, cells were washed with Hanks’ Balanced Salt Solution (HBSS; Thermo Fisher Scientific) and resuspended in cold BD Phosflow™ Perm Buffer III (BD Biosciences; Cat# 558050) for 30 mins at 4°C. After incubation, cells were washed with FACS buffer and incubated for 30 mins at room temperature with anti-pSMAD2/3 antibody (pS465/pS467; pS423/pS425; BD Biosciences; Cat# 562586). Post staining, T cells were centrifuged, resuspended in FACS buffer, and subsequently analyzed on a CytoFLEX flow cytometer (Beckman Coulter).

To demonstrate that TGFBR2 KO functionally abrogates TGF-β1 mediated suppression of T cell proliferation, engineered T cells were labelled with CellTrace^TM^ Violet and stimulated in plates pre-coated with immobilized anti-CD3 antibody (OKT3; Bio X Cell; Cat# BE0001-2) with or without exogenously added recombinant TGF-β1 (25 ng/mL). T cell proliferation was quantified by flow cytometry after 4 days of culture as described above.

### 6.13 OVCAR3 Tumor Model

OVCAR3-Luc2 cells were cultured for at least 3 passages before being trypsinized and resuspended in ice cold HBSS and Matrigel (Corning) in a 1:1 ratio, then injected subcutaneously into the dorsal hind flank (1.5×10^7^ cells/mouse). Animal health was monitored by clinical observations and body weight measurements. Tumor cell growth was monitored every 3-4 days using calipers or BioVolume (Fuel3D, UK) to track tumor burden. Once tumors reached target size, animals were randomized into treatment groups and injected into the tail vein with 100 µL of T cells suspended in HBSS (Corning) at the indicated cell dose. Tumor volume was calculated using the following formula when measured by calipers: Tumor volume = (length × width^2^)/2. For tumors measured with BioVolume, a machine learning algorithm using image segmentation of infrared and photographic images determined the length, width, and height of the tumor, and tumor volume was calculated as: Tumor volume = π × (length × width × height)/6. At indicated time points, animals were bled via the tail vein and approximately 50 µL of blood was collected into 96 well V bottom plates (Corning) containing 50µL of 0.5M EDTA (pH 8.0). For tumor rechallenge studies, treated mice that achieved tumor regression or treatment-naïve control mice were challenged with a bolus of 3×10^7^ OVCAR3 cells at a site adjacent to the primary tumor injection site. At the study endpoint, mice were euthanized, and spleens and blood were collected for downstream analysis. Spleen samples were mechanically dissociated into a single cell suspension using a GentleMACS Octo Dissociator (Miltenyi Biotec), passed through 70 µm cell strainer, and centrifuged. Red blood cell lysis was performed using ACK Lysis Buffer (Gibco), after which cells were washed and prepared for flow cytometry.

### 6.14 OCI-M1 Tumor Model

NOG mice were irradiated at 2 Gy and received DietGel and HydroGel (ClearH20) for nutritional and hydration support. Mice were then engrafted intravenously with 5×10^5^ OCI-M1-GFP-Luc2 tumor cells. At Day 10 post tumor engraftment, mice were randomized into treatment groups based on tumor burden, and 3×10^6^ or 1×10^7^ TCR+ T cells, suspended in HBSS (Corning), were administered intravenously. Animal health was monitored by clinical observation and body weight measurements. Tumor burden was assessed twice a week by IVIS bioluminescence imaging (PerkinElmer). Regions of Interest (ROIs) were defined and total flux values (photons/second) were calculated using Living Image Software.

### 6.15 T cell Pharmacokinetics by ddPCR

To quantify WT1-TCR T cells in mouse peripheral blood, genomic DNA (gDNA) was isolated from fresh blood samples using the Quick-DNA 96 Plus Kit (Zymo Research; Cat# D4071) following the manufacturer’s protocol and DNA concentration was quantified using a Stunner spectrophotometer (Unchained labs). The ddPCR primers and probes (Bio-Rad) designed to amplify the WT1-TCR comprised a forward primer targeting the TCRα constant region 5’ to the left homology arm of the HDRT and a reverse primer in the EF1α promoter region. The probe was complementary to a region within the homology arm of the HDRT and was FAM labelled. This primer/probe set was validated to specifically detect only the inserted transgenic WT1-TCR in human and mouse gDNA. A primer/probe set (HEX-labelled) targeting β-actin was included in each reaction as a reference for gDNA input. For ddPCR, known amounts of gDNA (50-200 ng) were mixed with 2x ddPCR Supermix for Probes (Bio-Rad; Cat# 1863024), Hind III HF restriction enzyme (New England biolabs; Cat# R3104S), forward and reverse primers (900nM each), and probes (250nM each). Droplets were generated using the Automated Droplet Generator (Bio-Rad) and samples were amplified using a thermocycler under the following cycling conditions: initial enzymatic activation for 10 min at 95°C; 50 cycles of denaturation (30 seconds at 94°C), annealing (1 minute at 60°C), and extension (4 minutes at 72°C); final enzymatic deactivation for 10 minutes at 98°C; followed by hold at 4°C. The temperature ramp rate all steps was 2°C/second. Droplet fluorescence was measured using a QX200 Droplet Reader (Bio-Rad), and WT1-TCR copy number (copies/µg gDNA) was calculated by normalizing copies per reaction to the input gDNA mass.

### 6.16 Statistical Analysis

Statistical analyses, including Student’s *t* test, one-way analysis of variance (ANOVA), and two-way ANOVA with post hoc multiple comparisons tests as indicated, were performed using GraphPad Prism software. Statistical significance was defined as *p* < 0.05. EC50 values were determined using a nonlinear regression model (least squares fit) with the 4-parameter dose-response equation of the GraphPad Prism software.

### 6.17 Data and Material Availability

Materials may be provided upon request at the sole discretion of Intellia Therapeutics, subject to completion of a material transfer agreement. All data supporting the findings of this study are presented in the main text or provided in the Supplementary Materials.

## 8 Acknowledgements

The authors would like to thank Mark Fetter, Long Chen, and the AAV core at Intellia for production and characterization of AAV in support of this study. Additionally, the authors would like to acknowledge Vishal Rakshe and Archana Swami for formulation and characterization of LNPs. Text here

## 9 Contributions

ID, JO, LC, AL, PS, IB, JP, IM, LM, JP, ES, DL, YK, JLC, JSM, designed and performed experiments and analyzed data. JB, QZ, EFL, LSL, ER, CB, BS, AP designed experiments, analyzed data, supervised the project, and contributed to the manuscript. ID, BS, and AP wrote the manuscript.

## 10 Competing Interests

ID, JO, LC, AL, PS, IB, JP, IM, JB, LM, JP, ES, DL, QZ, LSL, BS, and AP are all current or former employees of Intellia Therapeutics. AP., BS., have filed patent applications related to CD8 co-receptors. EFL has received research funding from Intellia Therapeutics. CB has been member of Advisory Board and Consultant for Intellia, TxCell, Novartis, GSK, Allogene, Kite/Gilead, Kiadis, Evir, Janssen, Genyo, Epsilen and received research support from Intellia Therapeutics. CB and ER are inventors on different patents on cancer immunotherapy and genetic engineering

## 12 Supplementary Figure Legends

**Figure S1. Enhancement of the cytotoxicity and cytokine secretion of CD4+ WT1-TCR+ T cells by expressing CD8αβ co-receptor (a)** Representative flow cytometric plots showing CD4+, CD4+CD8+, and CD8+ T cells following CRISPR-Cas9-mediated targeted integration of the CD8αβ co-receptor and WT1-TCR transgenes. **(b)** Enhanced cytotoxicity against 697 ALL tumor cells and **(c)** secretion of IFN-γ, TNF-α, GM-CSF, and Granzyme B in CD4+ WT1-TCR T cells following CD8αβ dual knock-in (DKI). Statistically significant but modest improvement of CD8+ WT1-TCR T cell activity was also observed following CD8αβ insertion. Symbols in (b) and bars in (c) represent mean ± SD (n = 3). **p<0.01, ****p<0.0001.

**Figure S2. Screening CD8 costimulatory fusion receptors with and without CD8 palmitoylation motifs (a)** Schematic of representative CD8-costimulatory chimera with re-inclusion of the CD8β chain palmitoylation domain (HLCCRR, Palm) or the palmitoylation domain with additional RR residues (HLCCRRRR, PalmR4). **(b)** Bar plots showing the transgene insertion rates (blue) and expression levels (MFI, orange) in CD4+ T cells after targeted integration of the indicated CD8 co-receptor design. **(c)** Serial rechallenge assay of WT1-TCR T cells with GFP+ OCI-M1 cells showed no improvement upon re-inclusion of Palm or PalmR4 motifs in in CD8 chimeric receptors containing 41BB (top) or modified CD28^47^ (bottom)signaling domains. # Denotes data points affected by an electrical infrastructure failure during the first ∼24h of the assay. **(d)** Schematic design of CD8 fusion constructs with 41BB ICD fused to the full length CD8α or CD8β intracellular sequences. (e) IFN-γ secretion was measured following co-culture of OVCAR3 tumor cells with WT1-TCR T cells carrying the indicated CD8 fusion constructs.

**Figure S3. Functional knockout of TGFBR2** (a) WT1-TCR T cells engineered with or without TGFBR2 KO were exposed to TGFβ1 and assessed for phosphorylated SMAD2/3 (pSMAD2/3) by flow cytometry. Data show abrogation of pSMAD2/3 induction in TGFBR2 KO cells and are representative of multiple experiments. (b) CTV labelled WT1-TCR T cells with or without TGFBR2 KO were stimulated with anti-CD3 antibody in the presence or absence of soluble TGFβ1. Proliferation was monitored by flow cytometry and quantified as the number of Vβ8+ (WT1-TCR) CTV^low^ cells/µL using counting beads. Data show prevention of TGFβ1-mediated suppression of T cell proliferation upon TGFBR2 KO. Bars represent mean ± SD (n=3, ****p<0.001). (c) OVCAR3 tumors were established subcutaneously (s.c.) and mice were treated with WT1-TCR T cells as described in Figure 5. WT1-TCR T cells with TGFBR2 KO alone were not effective in slowing tumor progression.

**Figure S4. Flow cytometric analysis of mouse spleens in the OVCAR3 rechallenge cohorts (a)** Flow cytometry was used to characterize WT1-TCR+ (Vβ8+), CD8+ WT1-TCR+, and CD4+CD8+ WT1-TCR+ T cells in the spleens of treated mice at the endpoint of the OVCAR3 rechallenge model (Figure 5). Each symbol represents an individual mouse. **(b)** Representative flow cytometric plots showing the gating of CD8+ stem cell memory T cells (Tscm: CD45RA+ CCR7+ CD28+ CD62L+).

**Figure S5. CD8-41BB receptors do not change the HLA-A*02:01 restriction or specificity profile of WT1-TCR cells** Cytotoxicity assays assessing the activity of WT1-TCR T cells against **(a)** WT1+K562 cells with or without HLA-A*02:01 expression or **(b)** WT1-HLA-A*02:01+ H441 cells with or without WT1-GFP expression confirmed that WT1-dependent, HLA-A*02:01-restricted killing is retained upon CD8-41BB co-receptor expression (n=2). **(c)** Quantification of live HLA-A*02:01+ WT1+ AML samples (n=2) and HLA-A*02:01-WT1+ AML sample (n=1) after co-culture with WT1-TCR T cells or WT1-TCR T cells co-expressing CD8-41BB co-receptor and TGFBR2 KO (E:T=1:12), normalized to the viability of TCR KO control cells, further confirming HLA-A*02:01-restricted activity. **(d)** Heatmap depicting the cytotoxicity of H441-GFP cells peptide-pulsed with the indicated WT1-VLD peptide, alanine-substitutedWT1 peptide variants, or the WT1-RMF negative control peptide. Shading represents mean % killing (n=3) of H441-GFP cells when pulsed at a peptide concentration approximating the EC90 determined for the WT1-TCR. **(e)** Quantification of IFN-γ secreted by WT1-TCR+ T cells expressing the indicated CD8 co-receptor following 24h co-culture with OVCAR3 cells or positively selected CD34+ bone marrow cells. Bars represent mean ± SD, (n=2). **(f)** Quantification of live HLA-A*02:01+ AML and HLA-A*02:01+ healthy donor bone marrow mononuclear cells (BMMCs post co-culture with WT1-TCR T cells alone or WT1-TCR T cells co-expressing CD8αβ or CD8-41BB co-receptor with TGFBR2 KO (E:T=1:12), demonstrating no additional cytotoxicity against healthy donor BMMCs upon expression of CD8αβ or CD8-41BB co-receptors.

## References

1. Cappell KM, Kochenderfer JN. Long-term outcomes following CAR T cell therapy: what we know so far. Nat Rev Clin Oncol. 2023;20(6):359–371. doi:10.1038/s41571-023-00754-1

2. Baulu E, Gardet C, Chuvin N, Depil S. TCR-engineered T cell therapy in solid tumors: State of the art and perspectives. Sci Adv. 2023;9(7):eadf3700. doi:10.1126/sciadv.adf3700

3. Borst J, Ahrends T, Bąbała N, Melief CJM, Kastenmüller W. CD4+ T cell help in cancer immunology and immunotherapy. Nat Rev Immunol. 2018;18(10):635–647. doi:10.1038/s41577-018-0044-0

4. Laugel B, Berg HA van den, Gostick E, et al. Different T Cell Receptor Affinity Thresholds and CD8 Coreceptor Dependence Govern Cytotoxic T Lymphocyte Activation and Tetramer Binding Properties*. J Biol Chem. 2007;282(33):23799–23810. doi:10.1074/jbc.m700976200

5. Yamamoto T, Hattori M, Yoshida T. Induction of T-cell activation or anergy determined by the combination of intensity and duration of T-cell receptor stimulation, and sequential induction in an individual cell. Immunology. 2007;121(3):383–391. doi:10.1111/j.1365-2567.2007.02586.x

6. Chow A, Perica K, Klebanoff CA, Wolchok JD. Clinical implications of T cell exhaustion for cancer immunotherapy. Nat Rev Clin Oncol. 2022;19(12):775–790. doi:10.1038/s41571-022-00689-z

7. Abate-Daga D, Hanada K ichi, Davis JL, Yang JC, Rosenberg SA, Morgan RA. Expression profiling of TCR-engineered T cells demonstrates overexpression of multiple inhibitory receptors in persisting lymphocytes. Blood. 2013;122(8):1399–1410. doi:10.1182/blood-2013-04-495531

8. Stegen SJC van der, Hamieh M, Sadelain M. The pharmacology of second-generation chimeric antigen receptors. Nat Rev Drug Discov. 2015;14(7):499–509. doi:10.1038/nrd4597

9. Anderson VE, Brilha SS, Weber AM, et al. Enhancing Efficacy of TCR-engineered CD4+ T Cells Via Coexpression of CD8α. J Immunother. 2023;46(4):132–144. doi:10.1097/cji.0000000000000456

10. Kessels HWHG, Schepers K, Boom MD van den, Topham DJ, Schumacher TNM. Generation of T Cell Help through a MHC Class I-Restricted TCR. J Immunol. 2006;177(2):976–982. doi:10.4049/jimmunol.177.2.976

11. Lahman MC, Schmitt TM, Paulson KG, et al. Targeting an alternate Wilms’ tumor antigen 1 peptide bypasses immunoproteasome dependency. Sci Transl Med. 2022;14(631):eabg8070. doi:10.1126/scitranslmed.abg8070

12. Liu X, Ranganathan R, Jiang S, et al. A Chimeric Switch-Receptor Targeting PD1 Augments the Efficacy of Second-Generation CAR T Cells in Advanced Solid Tumors. Cancer Res. 2016;76(6):1578–1590. doi:10.1158/0008-5472.can-15-2524

13. Sailer N, Fetzer I, Salvermoser M, et al. T-Cells Expressing a Highly Potent PRAME-Specific T-Cell Receptor in Combination with a Chimeric PD1-41BB Co-Stimulatory Receptor Show a Favorable Preclinical Safety Profile and Strong Anti-Tumor Reactivity. Cancers. 2022;14(8):1998. doi:10.3390/cancers14081998

14. Oda SK, Daman AW, Garcia NM, et al. A CD200R-CD28 fusion protein appropriates an inhibitory signal to enhance T-cell function and therapy of murine leukemia. Blood. 2017;130(22):2410–2419. doi:10.1182/blood-2017-04-777052

15. Anderson KG, Oda SK, Bates BM, et al. Engineering adoptive T cell therapy to co-opt Fas ligand-mediated death signaling in ovarian cancer enhances therapeutic efficacy. J Immunother Cancer. 2022;10(3):e003959. doi:10.1136/jitc-2021-003959

16. Wang X, Teng F, Kong L, Yu J. PD-L1 expression in human cancers and its association with clinical outcomes. OncoTargets Ther. 2016;9:5023–5039. doi:10.2147/ott.s105862

17. Mann B, Gratchev A, Böhm C, et al. FasL is more frequently expressed in liver metastases of colorectal cancer than in matched primary carcinomas. Br J Cancer. 1999;79(7-8):1262–1269. doi:10.1038/sj.bjc.6690202

18. Noviello M, Manfredi F, Ruggiero E, et al. Bone marrow central memory and memory stem T-cell exhaustion in AML patients relapsing after HSCT. Nat Commun. 2019;10(1):1065. doi:10.1038/s41467-019-08871-1

19. Verdun N, Marks P. Secondary Cancers after Chimeric Antigen Receptor T-Cell Therapy. N Engl J Med. Published online 2024. doi:10.1056/nejmp2400209

20. Ruggiero E, Carnevale E, Prodeus A, et al. CRISPR-based gene disruption and integration of high-avidity, WT1-specific T cell receptors improve antitumor T cell function. Sci Transl Med. 2022;14(631):eabg8027. doi:10.1126/scitranslmed.abg8027

21. Jaigirdar A, Rosenberg SA, Parkhurst M. A High-avidity WT1-reactive T-Cell Receptor Mediates Recognition of Peptide and Processed Antigen but not Naturally Occurring WT1-positive Tumor Cells. J Immunother. 2016;39(3):105–116. doi:10.1097/cji.0000000000000116

22. Qi X wei, Zhang F, Wu H, et al. Wilms’ tumor 1 (WT1) expression and prognosis in solid cancer patients: a systematic review and meta-analysis. Sci Rep. 2015;5(1):8924. doi:10.1038/srep08924

23. Meng K, Cao J, Dong Y, Zhang M, Ji C, Wang X. Application of Bioinformatics Analysis to Identify Important Pathways and Hub Genes in Ovarian Cancer Affected by WT1. Front Bioeng Biotechnol. 2021;9:741051. doi:10.3389/fbioe.2021.741051

24. Tan MP, Dolton GM, Gerry AB, et al. Human leucocyte antigen class I-redirected anti-tumour CD4+ T cells require a higher T cell receptor binding affinity for optimal activity than CD8+ T cells. Clin Exp Immunol. 2016;187(1):124–137. doi:10.1111/cei.12828

25. Hockemeyer D, Soldner F, Beard C, et al. Efficient targeting of expressed and silent genes in human ESCs and iPSCs using zinc-finger nucleases. Nat Biotechnol. 2009;27(9):851–857. doi:10.1038/nbt.1562

26. Hayashi H, Kubo Y, Izumida M, Matsuyama T. Efficient viral delivery of Cas9 into human safe harbor. Sci Rep. 2020;10(1):21474. doi:10.1038/s41598-020-78450-8

27. Arcaro A, Grégoire C, Boucheron N, et al. Essential Role of CD8 Palmitoylation in CD8 Coreceptor Function. J Immunol. 2000;165(4):2068–2076. doi:10.4049/jimmunol.165.4.2068

28. Pang DJ, Hayday AC, Bijlmakers MJ. CD8 Raft Localization Is Induced by Its Assembly into CD8αβ Heterodimers, Not CD8αα Homodimers*. J Biol Chem. 2007;282(18):13884–13894. doi:10.1074/jbc.m701027200

29. Knezevic L, Wachsmann TLA, Francis O, et al. High-affinity CD8 variants enhance the sensitivity of pMHCI antigen recognition via low-affinity TCRs. J Biol Chem. 2023;299(8):104981. doi:10.1016/j.jbc.2023.104981

30. Nam KO, Shin SM, Lee HW. Cross-linking of 4-1BB up-regulates IL-13 expression in CD8+ T lymphocytes. Cytokine. 2006;33(2):87–94. doi:10.1016/j.cyto.2005.12.003

31. Kochenderfer JN, Somerville RPT, Lu T, et al. Lymphoma Remissions Caused by Anti-CD19 Chimeric Antigen Receptor T Cells Are Associated With High Serum Interleukin-15 Levels. J Clin Oncol. 2017;35(16):JCO.2016.71.302. doi:10.1200/jco.2016.71.3024

32. Canale FP, Ramello MC, Núñez N, et al. CD39 expression defines cell exhaustion in tumor-infiltrating CD8+ T cells. Cancer Res. 2017;78(1):canres.2684.2016. doi:10.1158/0008-5472.can-16-2684

33. Amerongen RA van, Hagedoorn RS, Remst DFG, et al. WT1-specific TCRs directed against newly identified peptides install antitumor reactivity against acute myeloid leukemia and ovarian carcinoma. J Immunother Cancer. 2022;10(6):e004409. doi:10.1136/jitc-2021-004409

34. Roth TL, Puig-Saus C, Yu R, et al. Reprogramming human T cell function and specificity with non-viral genome targeting. Nature. 2018;559(7714):405–409. doi:10.1038/s41586-018-0326-5

35. Wermke M, Tsimberidou AM, Mohamed A, et al. 959 Safety and anti-tumor activity of TCR-engineered autologous, PRAME-directed T cells across multiple advanced solid cancers at low doses – clinical update on the ACTengine® IMA203 trial. J Immunother Cancer. 2021;9(Suppl 2):A1009–A1009. doi:10.1136/jitc-2021-sitc2021.959

36. Wermke M, Alsdorf W, Araujo D, et al. Abstract PR018: IMA203 TCR-T targeting PRAME demonstrates potent anti-tumor activity in patients with different types of metastatic solid tumors. Mol Cancer Ther. 2023;22(12_Supplement):PR018–PR018. doi:10.1158/1535-7163.targ-23-pr018

37. Maher J, Brentjens RJ, Gunset G, Rivière I, Sadelain M. Human T-lymphocyte cytotoxicity and proliferation directed by a single chimeric TCRζ /CD28 receptor. Nat Biotechnol. 2002;20(1):70–75. doi:10.1038/nbt0102-70

38. Krause A, Guo HF, Latouche JB, Tan C, Cheung NKV, Sadelain M. Antigen-dependent CD28 Signaling Selectively Enhances Survival and Proliferation in Genetically Modified Activated Human Primary T Lymphocytes. J Exp Med. 1998;188(4):619–626. doi:10.1084/jem.188.4.619

39. Milone MC, Fish JD, Carpenito C, et al. Chimeric Receptors Containing CD137 Signal Transduction Domains Mediate Enhanced Survival of T Cells and Increased Antileukemic Efficacy In Vivo. Mol Ther. 2009;17(8):1453–1464. doi:10.1038/mt.2009.83

40. Ostroumov D, Fekete-Drimusz N, Saborowski M, Kühnel F, Woller N. CD4 and CD8 T lymphocyte interplay in controlling tumor growth. Cell Mol Life Sci. 2018;75(4):689–713. doi:10.1007/s00018-017-2686-7

41. Zhang S, Tang TH, Kinsella S, et al. A CD8αβ co-receptor modified to contain an intracellular CD28 signaling tail enhances TCR-engineered T cell function independent of solid-tumor-associated co-stimulatory ligands. Nat Commun. 2026;17(1):753. doi:10.1038/s41467-025-67446-5

42. Kalaitsidou M, Moon OR, Sykorova M, et al. Signaling via a CD28/CD40 chimeric costimulatory antigen receptor (CoStAR^TM^), targeting folate receptor alpha, enhances T cell activity and augments tumor reactivity of tumor infiltrating lymphocytes. Front Immunol. 2023;14:1256491. doi:10.3389/fimmu.2023.1256491

43. Omer B, Cardenas MG, Pfeiffer T, et al. A Costimulatory CAR Improves TCR-based Cancer Immunotherapy. Cancer Immunol Res. 2022;10(4):512–524. doi:10.1158/2326-6066.cir-21-0307

44. Dobrin A, Lindenbergh PL, Shi Y, et al. Synthetic dual co-stimulation increases the potency of HIT and TCR-targeted cell therapies. Nat Cancer. 2024;5(5):760–773. doi:10.1038/s43018-024-00744-x

45. Klobuch S, Seijkens TTP, Schumacher TN, Haanen JBAG. Tumour-infiltrating lymphocyte therapy for patients with advanced-stage melanoma. Nat Rev Clin Oncol. 2024;21(3):173–184. doi:10.1038/s41571-023-00848-w

46. Wang Z, Ahmed S, Labib M, et al. Isolation of tumour-reactive lymphocytes from peripheral blood via microfluidic immunomagnetic cell sorting. Nat Biomed Eng. 2023;7(9):1188–1203. doi:10.1038/s41551-023-01023-3

47. Nguyen P, Moisini I, Geiger TL. Identification of a murine CD28 dileucine motif that suppresses single-chain chimeric T-cell receptor expression and function. Blood. 2003;102(13):4320–4325. doi:10.1182/blood-2003-04-1255

48. Jetley U, Balwani I, Sharma P, et al. A differentiated and durable allogeneic strategy applicable to cell therapies. Cytotherapy. 2025;28(3):101991. doi:10.1016/j.jcyt.2025.10.001

49. Vavassori V, Ferrari S, Beretta S, et al. Lipid nanoparticles allow efficient and harmless ex vivo gene editing of human hematopoietic cells. Blood. 2023;142(9):812–826. doi:10.1182/blood.2022019333

