## Supplementary Figures for "Engineering antigen-driven co-stimulation and T helper cell activity into TCR-T cells with CD8-41BB fusion receptors enhances anti-tumor activity"

Supplementary Figure 1

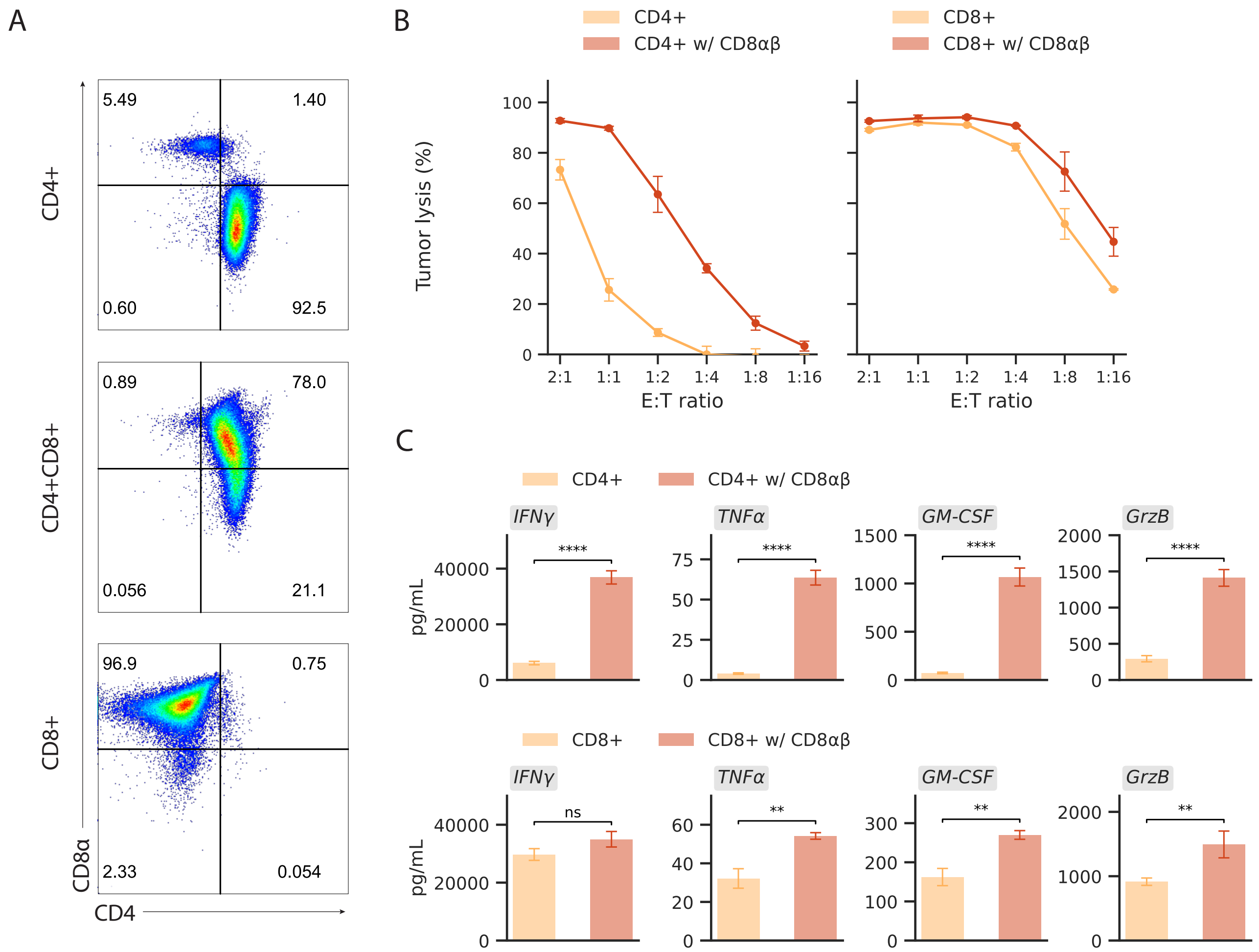

Supplementary Figure 2

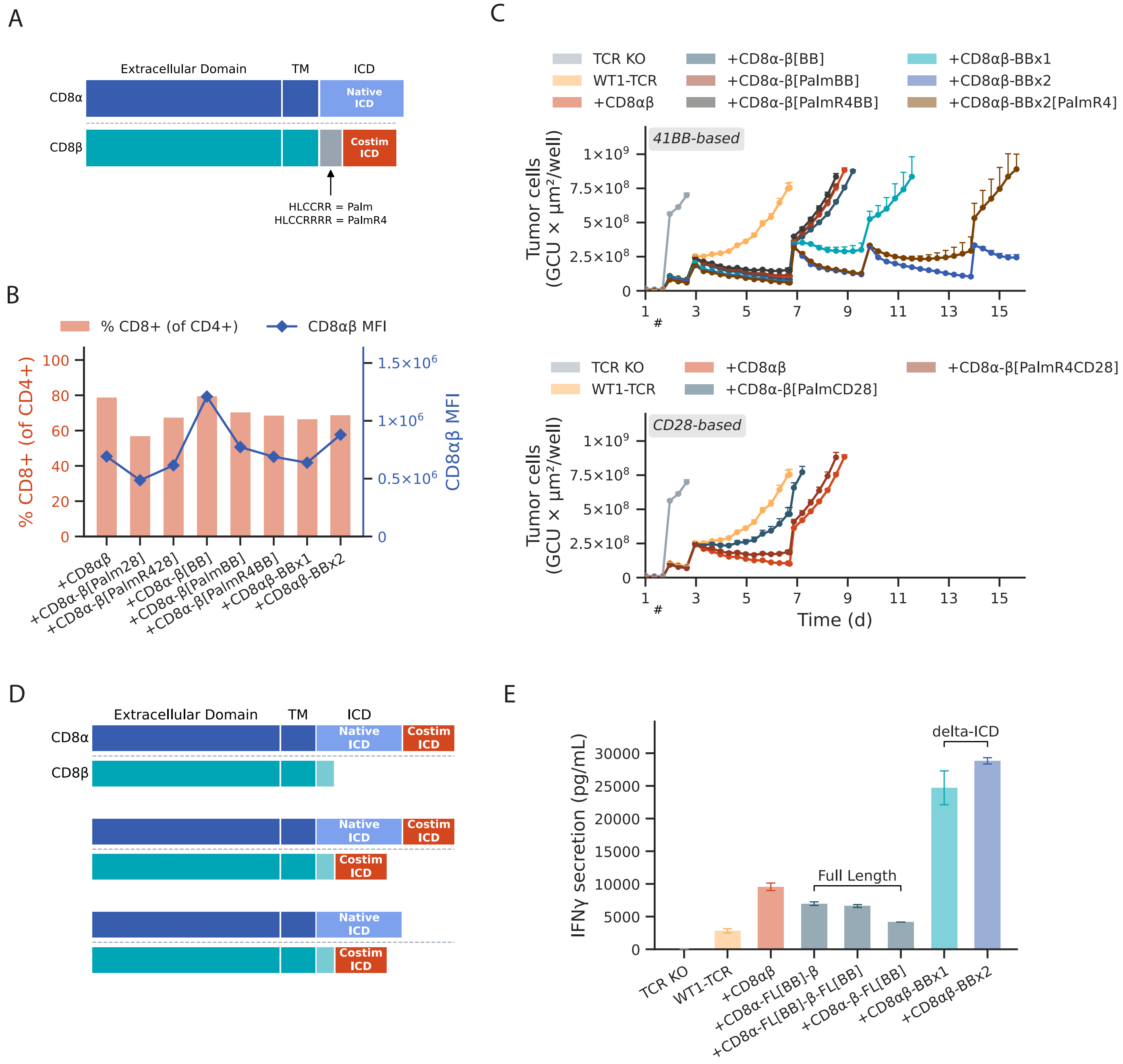

Supplementary Figure 3

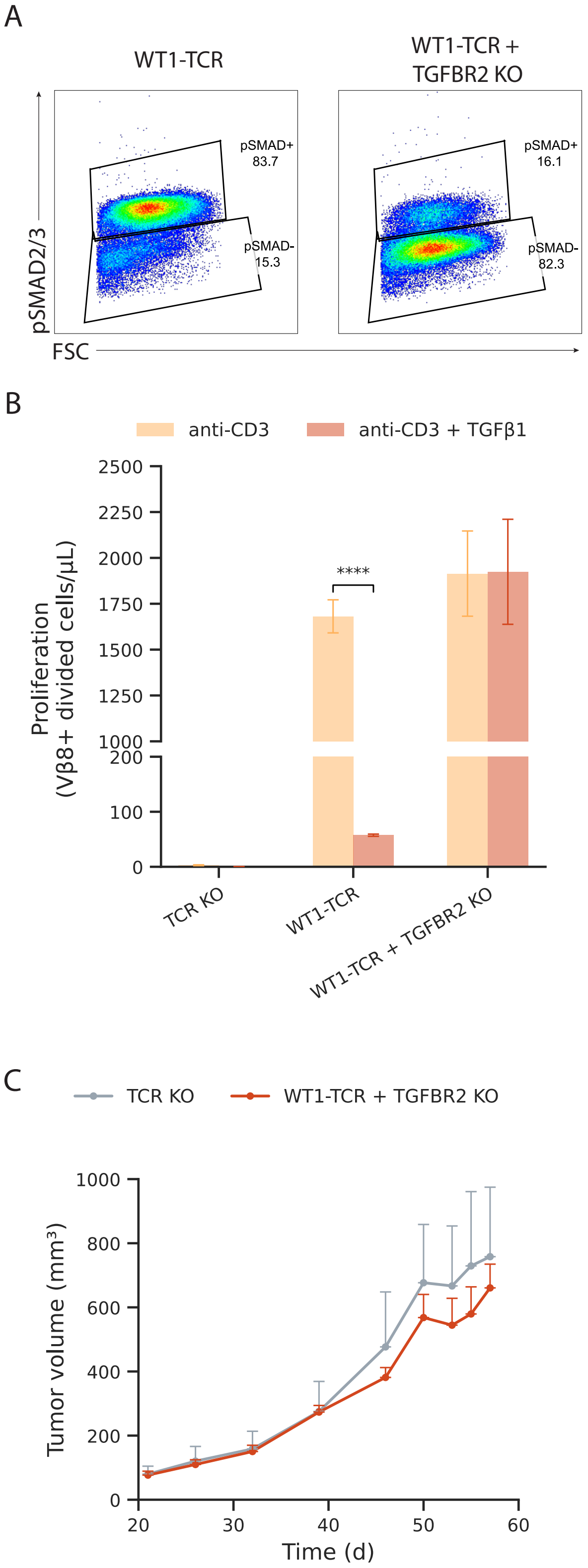

Supplementary Figure 4

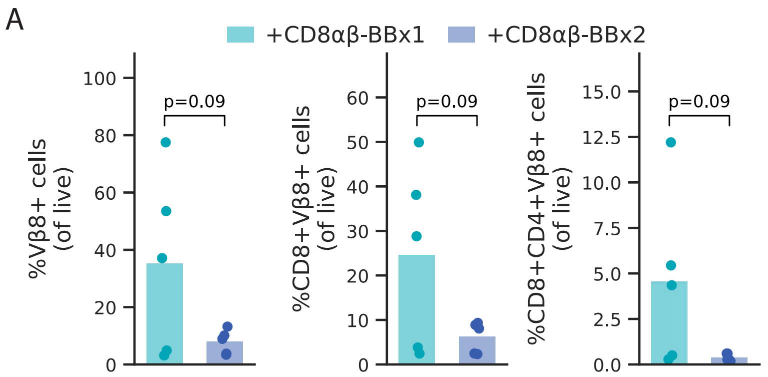


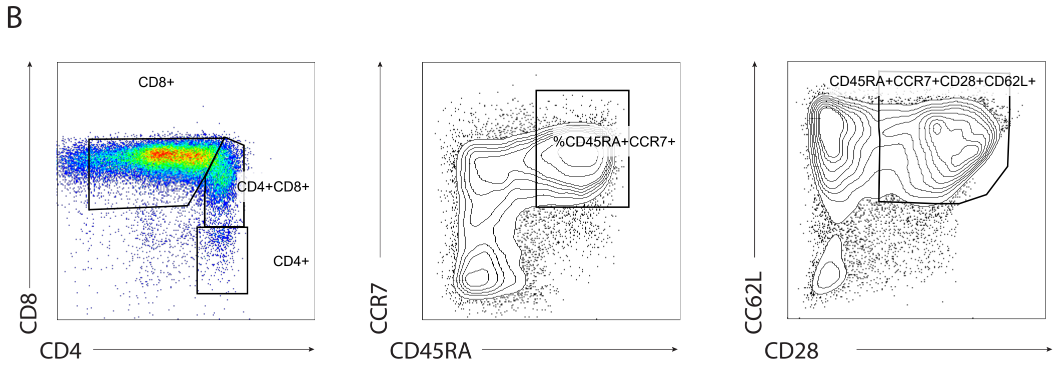

Supplementary Figure 5

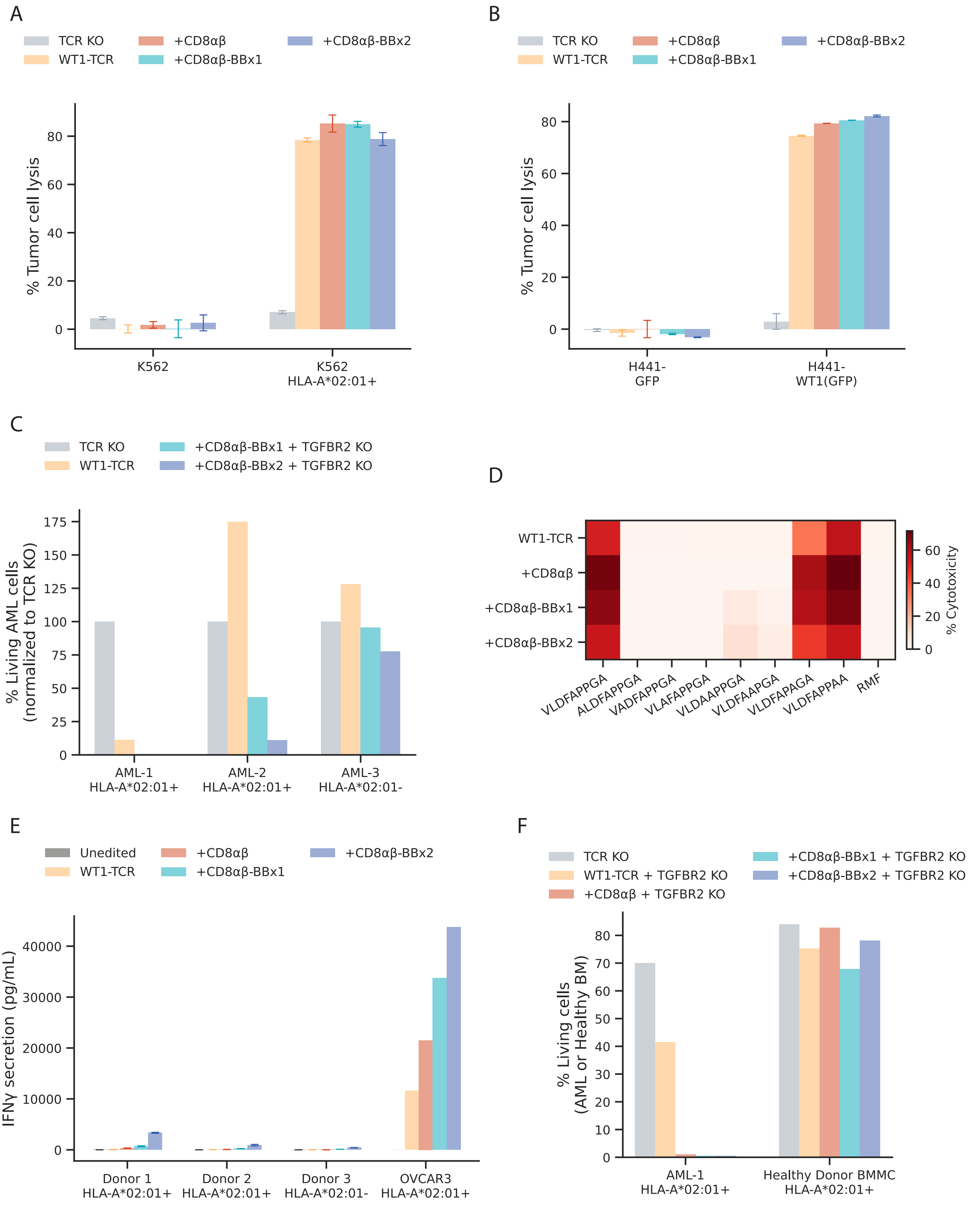
