## Supplementary Tables for "Engineering antigen-driven co-stimulation and T helper cell activity into TCR-T cells with CD8-41BB fusion receptors enhances anti-tumor activity"

**Table S1. Antibodies used for flow cytometry analysis.**

| **Antigen** | **Clone** | **Fluorophore** | **Cat#** | **Vendor** | **Host** | **Application** | **Dilution** |
| --- | --- | --- | --- | --- | --- | --- | --- |
| CD123 | 6H6 | BV605 | 306026 | BioLegend | Mouse | Surface | 1/200 |
| CD19 | HIB19 | PE | 302208 | BioLegend | Mouse | Surface | 1/200 |
| CD197 (CCR7) | G043H7 | BV421 | 353208 | BioLegend | Mouse | Surface | 1/200 |
| CD197 (CCR7) | G043H7 | APC | 353214 | BioLegend | Mouse | Surface | 1/200 |
| CD223 (LAG3) | 11C3C65 | BV510 | 369318 | BioLegend | Mouse | Surface | 1/100 |
| CD223 (LAG3) | 11C3C65 | AF488 | 369326 | BioLegend | Mouse | Surface | 1/100 |
| CD27 | M-T271 | PE | 356406 | BioLegend | Mouse | Surface |  |
| CD28 (CD28.2) | CD28.2 | NovaFluor Red 710 | H008T03R04 | Thermo Fisher | Mouse | Surface | 1/40 |
| CD3 | UCHT1 | BUV395 | 563548 | BD Biosciences | Mouse | Surface | 1/200 |
| CD3 | OKT3 | PerCP-Cy5.5 | 317336 | BioLegend | Mouse | Surface | 1/200 |
| CD3 | UCHT1 | V450 | 652356 | BD Biosciences | Mouse | Surface | 1/200 |
| CD33 | WM53 | BV711 | 303424 | BioLegend | Mouse | Surface | 1/200 |
| CD34 | 561 | BV785 | 343626 | BioLegend | Mouse | Surface | 1/200 |
| CD366 (Tim-3) | F38-2E2 | BV510 | 345030 | BioLegend | Mouse | Surface | 1/100 |
| CD38 | HB-7 | PerCP | 356622 | BioLegend | Mouse | Surface | 1/200 |
| CD39 | A1 | BV785 | 328240 | BioLegend | Mouse | Surface | 1/50 |
| CD4 | OKT4 | BV421 | 317434 | BioLegend | Mouse | Surface | 1/200 |
| CD4 | SK3 | BUV661 | 612962 | BD Biosciences | Mouse | Surface | 1/200 |
| CD4 | OKT4 | BV605 | 317438 | BioLegend | Mouse | Surface | 1/200 |
| CD45 | HI30 | PE-Cy5 | 304010 | BioLegend | Mouse | Surface | 1/200 |
| CD45 | 2B11 & HI30 | PerCP-Cy5.5 | 304008 | BioLegend | Mouse | Surface | 1/200 |
| CD45RA | HI100 | PE-Cy7 | 304126 | BioLegend | Mouse | Surface | 1/100 |
| CD45RA | HI100 | FITC | 304106 | BioLegend | Mouse | Surface | 1/200 |
| CD45RO | UCHL1 | PE-Dazzle594 | 304248 | BioLegend | Mouse | Surface | 1/100 |
| CD45RO | UCHL1 | PE/Cyanine7 | 304230 | BioLegend | Mouse | Surface | 1/200 |
| CD62L | DREG-56 | APC | 304810 | BioLegend | Mouse | Surface | 1/100 |
| CD62L | DREG-56 | APC-700 | 304820 | BioLegend | Mouse | Surface | 1/200 |
| CD69 | FN50 | APC/Cyanine7 | 310914 | BioLegend | Mouse | Surface | 1/200 |
| CD8 | SK1 | APC-Fire750 | 344746 | BioLegend | Mouse | Surface | 1/100 |
| CD8 | SK1 | PerCP-Cy5.5 | 344710 | BioLegend | Mouse | Surface | 1/200 |
| CD8 | SK1 | APC-700 | 344724 | BioLegend | Mouse | Surface | 1/200 |
| CD8α | RPA-T8 | BV785 | 301046 | BioLegend | Mouse | Surface |  |
| CD8β | QA20A40 | APC | 376706 | BioLegend | Mouse | Surface | 1/200 |
| CD8β | 2ST8.5H7 | BV421 | 568373 | BD Biosciences | Mouse | Surface | 1/200 |
| CD95 (Fas) | DX2 | BV650 | 305642 | BioLegend | Mouse | Surface | 1/200 |
| CellTrace™ Violet Cell Proliferation Kit (CTV) | N/A | BV450 | C34557 | Thermo Fisher | N/A | Cell Proliferation Tracking | 1/1000 |
| HLA-A2 | BB7.2 | APC | 343308 | BioLegend | Mouse | Surface | 1/200 |
| HLA-ABC | W6/32 | APC-Cy7 | 311426 | BioLegend | Mouse | Surface | 1/200 |
| HLA-DR | L243 | BV570 | 307638 | BioLegend | Mouse | Surface | 1/200 |
| IFNγ | B27 | APC | 506510 | BioLegend | Mouse | Intracellular | 1/100 |
| LAG3 | 11C3C65 | PE-Cy7 | 369310 | BioLegend | Mouse | Surface | 1/200 |
| LIVE/DEAD™ Fixable Violet Dead Cell Stain | N/A | Violet | L34955 | Thermo Fisher | N/A | Viability | 1/100 |
| PD-1 | NAT105 | BV605 | 367426 | BioLegend | Mouse | Surface | 1/50 |
| PD-1 | EH12.2H7 | AF700 | 329952 | BioLegend | Mouse | Surface | 1/200 |
| PD-L1 | 29E.2A3 | BV650 | 329740 | BioLegend | Mouse | Surface | 1/200 |
| PD-L2 | 24F.10C12 | PE-Dazzle594 | 329622 | BioLegend | Mouse | Surface | 1/200 |
| Smad2 (pS465/pS467)/Smad3 (pS423/pS425) | O72-670 | PE | 562586 | BD Biosciences | Mouse | Intracellular (phospho-flow) | 1/200 |
| TCR Vβ8 | JR2 (JR.2) | PE | 348104 | BioLegend | Mouse | Surface | 1/100 |
| TCR Vβ8 | JR2 (JR.2) | FITC | 555606 | BD Biosciences | Mouse | Surface | 1/200 |
| TIM3 | F38-2E2 | PE-Dazzle594 | 345034 | BioLegend | Mouse | Surface | 1/200 |
| TNF-α | MAb11 | BV650 | 502938 | BioLegend | Mouse | Intracellular | 1/100 |
| ViaKrome 808 Fixable Viability Dye | -- | ViaKrome 808 | C36628 | Beckman Coulter | N/A | Viability | 1/100 |
| WT1 | WT1/3477R | AF700 | NBP3-08966AF700 | R&D Systems | Rabbit | Intranuclear | 1/50 |
| WT1 Dextramer (HLA-A*0201/VLDFAPPGA)* | N/A | PE | WB03469 | Immudex | Recombinant MHC-peptide | Surface — Antigen-specific T cell detection | 1/20 |
| Zombie Aqua™ Fixable Viability Kit | N/A | BV510 | 423101 | BioLegend | N/A | Viability | 1/300 |
| Zombie Yellow™ Fixable Viability Kit | -- | BV525 | 423104 | BioLegend | N/A | Viability | 1/100 |

** WT1 Dextramer (HLA-A*0201/VLDFAPPGA): RUO; custom order, peptide VLDFAPPGA, allele HLA-A*0201. Cat# WB03469 is representative of (HLA-A*0201/VLDFAPPGA)/no fluorophore.*
